# Macrophage Circadian Rhythms are Differentially Affected Based on Stimuli

**DOI:** 10.1101/2021.07.01.450771

**Authors:** Sujeewa S. Lellupitiyage Don, Javier A. Mas-Rosario, Hui-Hsien Lin, Minh N. Nguyen, Stephanie R. Taylor, Michelle E. Farkas

## Abstract

Macrophages are white blood cells of the innate immune system that play disparate roles in homeostasis and immune responses. As a result, they have the capability to alter their phenotypes to pro-inflammatory (M1) or anti-inflammatory (M2) subtypes in response to their environment. 8-15% of the macrophage transcriptome has circadian oscillations, including genes closely related to their functioning. As circadian rhythms are also associated with cellular phenotypes, we hypothesized that the polarization of macrophages to opposing subtypes might differently affect their circadian rhythms. We tested this by tracking the circadian rhythms of the mouse model macrophage cell line, RAW 264.7, which was stably transfected with *Bmal1:luc* and *Per2:luc* reporters, representing a positive and a negative component of the core molecular clock. The strength of rhythmicity was assessed using three measures: the relative power of the circadian band in the power spectral density, the rhythmicity index computed as the height of the third peak of the correlogram, and the maximum value of the chi square periodorgram. The period and amplitudes of the de-trended, smoothed time-series were estimated both by fitting to a damped cosine curve and by identifying the peak and trough of each cycle. M1 polarization decreased amplitudes and rhythmicities of both *Bmal1:luc* and *Per2:luc*, but did not significantly affect periods, while M2 polarization increased periods and caused no substantial alterations to amplitudes or rhythmicity. As macrophage phenotypes are also altered in the presence of cancer cells, we then tested the circadian effects of conditioned media from two mouse breast cancer cell lines on macrophage circadian rhythms. Media from highly agressive 4T1 cells caused loss of rhythmicity, while media from less aggressive EMT6 cells yielded no changes. As macrophages are known to play roles in tumors, and oncogenic features are associated with circadian rhythms, we also tested whether conditioned media from macrophages can alter circadian rhythms of cancer cells. We found that conditioned media from RAW 264.7 cells resulted in lower circadian rhythmicities and periods, but higher circadian amplitudes in human osteosarcoma, U2OS-*Per2:luc* cells. Taken together, our study shows that different circadian characteristics exist based on macrophage phenotypes, and suggests further that there is an association between circadian rhythms and macrophage polarization state. Additionally, our data shows that macrophages treated with breast cancer-conditioned media have circadian phenotypes similar to those of the M1 subtype, and cancer cells treated with macrophage-conditioned media have circadian alterations, providing insight to another level of cross-talk between macrophages and cancer cells that merits further investigation.

## Introduction

The immune system is a network of cells, organs, and biomolecular entities that protects organisms against foreign pathogens. Multiple immune-related activities are regulated in a circadian manner, meaning that they occur with an approximately twenty-four hour cycle. These include natural killer (NK) cell cytokine production and immunity against pathogens [1], numbers of particular white blood cell types (e.g., hematopoietic stem cells, B cells, and T cells) in the blood [2], and/or lymph nodes [3], migration of hematopoietic cells, neutrophils, and monocytes to tissues [2], phagocytic activity of neutrophils, [4] and expression of cytolytic factors by NK cells [1,5]. As a result, circadian time can determine immune responses to stimuli. For example, sensitivity to lipopolysaccharide (LPS, an endotoxin found in outer membranes of gram negative bacteria) [6] and inflammatory responses against *Salmonella enterica serovar Typhimurium* [7] have been found to be circadian, with increased responses at the start of the active phase and early rest period of mice, respectively. Mice infected with *Leishmania* [8] or *Trichuris muris* [9] pathogens in the morning showed lower parasite loads relative to those infected in the evening. Mouse models of peritoneal inflammation show more recruitment of inflammatory monocytes to the peritoneal cavity, spleen, and liver in the afternoon versus the morning [10], while in mouse models of myocardial infarction, neutrophil recruitment to cardiac tissue peaks in the evening [11]. In humans, which are diurnal (as opposed to mice, which are nocturnal), immune responses can also vary in a circadian manner. Allergic symptoms are increased between midnight and the early morning [11,12]; influenza vaccinations in the morning result in increased antibody concentrations compared to those administered in the afternoon [13], and asthma worsened lung function and resulted in increased sputnum leukocytes and eosinophils in the early morning, compared to in the late afternoon [14].

Macrophages are phagocytic white blood cells involved not only in the innate immune response, but also development, homeostasis, and tissue repair [15]. Macrophages shift their functional state to respond to physiological challenges or environmental factors. Specific characteristics and roles are associated with their polarization to subtypes broadly classified as classically (M1) or alternatively (M2) activated, which are inflammatory or immune-suppressing, respectively [15,16]. Macrophages have been shown to possess their own intrinsic clocks that operate autonomously [17], with a functional core molecular circadian clock (comprised of a negative feedback loop with BMAL1 and CLOCK proteins as activators, and PER and CRY proteins as repressors) [18]. Approximately 8-15% of the macrophage transcriptome has circadian oscillations, including genes required for pathogen recognition and responses [17,19]. Furthermore, multiple macrophage functions are circadian-dependent, such as recruitment to infected tissue [8], generation of chemokines and cytokines, and phagocytosis [20,21]. Considering that macrophages have highly divergent functions, from tissue remodeling [22] and angiogenesis [23] to generation of reactive oxygen species (ROS) [24] and phagocytosis [20,21], it is important to address how circadian oscillations might change concomitantly with phenotype.

Previous studies have indicated that there is a link between macrophage activation and circadian rhythms and/or molecular clock elements. Previous studies evaluated the impacts of LPS on circadian clocks in primary, peritoneal macrophages derived from mPER2:luc mice [25] and LPS, LPS/IFN-γ and IL-4 stimulation *mPer2:luc* mouse bone marrow-derived macrophages (BMDMs) [26] via time-dependent luminometry and RT-PCR assessments, finding that immune-activating treatments affected circadian period, phase, and amplitude. Curtis et al. also observed LPS-mediated effects, and assessed additional characteristics in BMDMs and peritoneal macrophages, and isolated human macrophages, at two individual time points. In this instance, *Per2* mRNA and protein expression were significantly increased, while those of *Bmal1* decreased [27]. In separate work, downregulation of *Bmal1* in mouse BMDMs increased the expression of proinflammatory cytokines, which suggested that *Bmal1* may act as an anti-inflammatory molecule in macrophages [28].

Associated with *Bmal1*, Curtis et al. also found that LPS significantly decreased levels of its transcriptional reporessor, *REV-ERBα*, but did not affect the presence of its transcriptional activator, *RORα* [27]. However, another study showed that stimulation with a combination of LPS and IFN-γ, which polarizes macrophages to the pro-inflammatory M1 subtype, increased *REV-ERBα* mRNA and protein levels in differentiated THP-1 macrophages. Concurrently, treatment with IL-4, which polarizes macrophages to the anti-inflammatory M2 subtype, resulted in REV-ERBα reduction [29]. These studies suggest that pro- and anti-inflammatory polarizations can differently affect circadian clock gene expression.

However, the implications of macrophage stimulation on circadian oscillations are largely unknown. Because the circadian clock is a dynamic system, it requires continuous monitoring to obtain sufficiently high resolution data for the accurate characterization of its patterns. For this reason, we have applied the use of luciferase reporters for real-time tracking of circadian oscillations in macrophages to study the relationship between macrophage activation and the biological clock.

On account of their distinct and opposing functions, and the results of prior studies, we hypothesized that polarization of macrophages to pro- and anti-inflammatory subtypes would affect their circadian rhythms differently. To study this, we generated stable reporter cell lines assessing a positive (*Bmal1*) and a negative (*Per2*) component of the core circadian clock, using RAW 264.7 macrophages. We tracked their oscillations via real-time luminometry following polarization to M1 and M2 subtypes using standard cytokine treatments (M1 via LPS or a combination of LPS/IFN-γ, M2 via IL-4). Polarization states of macrophages were verified by evaluating levels of M1- and M2-associated markers via RT-PCR. M1 polarization resulted in decreased amplitudes and rhythmicities of both *Bmal1:luc* and *Per2:luc* reporters, but did not significantly affect periods. On the other hand, M2 polarization resulted in increased periods, but not significant alterations to amplitudes or rhythmicity.

Because macrophages play major roles in cancers, which can also affect their phenotypes, we also assessed the effects of cancer cell-conditioned media on macrophage oscillations. Exposure to media from 4T1 cells, generally considered highly aggressive, resulted in significant loss of rhythmicity, while media from EMT6 (less aggressive) cells yielded no detectable changes. Finally, we sought to determine whether macrophages could affect circadian rhythms of cancer cells. Following treatment with RAW 264.7-conditioned media, human osteosarcoma cells (U2OS-*Per2:luc*) showed reduced circadian rhythmicity and period, and enhanced circadian amplitudes. Taken together, our results suggest that macrophage polarization, and therefore function, is linked to circadian oscillations, and changes therein.

## Materials and Methods

### Cell Culture

RAW 264.7 murine macrophage, and 4T1 and EMT6 murine mammary carcinoma cells were obtained from the ATCC; 293T human embryonic kidney cells were obtained from the Jerry Laboratory (Dept. of Veterinary and Animal Sciences, UMass Amherst); U2OS human osteosarcoma cells were obtained from the Wadsworth laboratory (Dept. of Biology, UMass Amherst). RAW 264.7, EMT6, and 4T1 cells were maintained in high glucose DMEM (Gibco), supplemented with 10% Fetal Bovine Serum (FBS; Corning), 1% penicillin-streptomycin (Gibco), and 1% L-glutamine (Gibco). 293T cells were maintained in DMEM/F12 media (Gibco), supplemented with 10% FBS, 1% penicillin-streptomycin, and 0.015 mg/mL gentamicin (Gibco). U2OS cells were maintained in DMEM, supplemented with 10% FBS, 1% penicillin-streptomycin, 1% L-Glutamine, 1% non-essential amino acids (NEAA; HyClone), and 1% sodium pyruvate (Gibco). Experiment-specific media preparations are indicated below. All cells were incubated at 37 °C under 5% CO_2_ atmosphere, unless otherwise noted.

### Lentiviral Transfections

The generation of *Bmal1:luc* and *Per2:luc* plasmids have been described previously [30], as has their transfection into U2OS cells [31]. Subsequent stable transfections of these reporters into RAW 264.7 cells were performed similarly to as in other studies [31,32]. Briefly, 3 x 10^6^ 293T cells were seeded in 60 mm culture dishes and transiently transfected with 3 µg psPAX2 packaging plasmid, 2 µg pMD2.G envelope plasmid, and 5 µg *Bmal1:luc* or *Per2:luc* plasmid constructs using Lipofectamine3000 (Thermo Fisher Scientific). After 48 h incubation, lentiviral particles were harvested and filtered through a 0.45 µm membrane (Thermo Fisher Scientific). 9 mL lentivirus-containing supernatant was combined with 9 mL RAW 264.7 growth media containing 10 µg/mL polybrene (Sigma). Cells were seeded in T25 culture flasks at 2 x 10^5^ cells/mL and incubated under standard conditions until 70-80% confluence was reached.

At this stage, culture media was removed and 6 mL lentivirus-containing media was added. After 2 days of infection, media was replaced with selection media (DMEM with all growth supplements plus 4 µg/mL puromycin (Gibco)). The selection media was changed once every 3 days for 4 weeks.

### Cell Synchronization

For RAW 264.7 cell synchronization, cells were seeded in 35 mm culture dishes at 2 x 10^5^ cells/mL and incubated for approximately 24 h. Media was then aspirated, cells washed with phosphate-buffered saline (PBS; Gibco), and synchronized by subjecting to starvation conditions by adding starvation media (L-15 media (Gibco) with 1% penicillin-streptomycin and 1% L-glutamine) and incubating for 18 h. Starvation media was removed and replaced with L-15 media containing only 100 nM dexamethasone (Sigma-Aldrich) for 2 h. Both starvation and synchronization were carried out at 37 °C under 5% CO_2_ atmosphere. For U2OS cell synchronization, cells were seeded in 35 mm culture dishes at 2 x 10^5^ cells/mL and incubated for approximately 24 h. Media was then aspirated and replaced with U2OS culture media containing 100 nM dexamethasone, followed by incubation for 2 h at 37 °C under 5% CO_2_ atmosphere.

### Cell Treatments with Cytokines and Conditioned Media

Lipopolysaccharide (LPS; Sigma-Aldrich), IFN-γ (BD Biosciences), and IL-4 (BioLegend) were prepared in PBS at concentrations of 1000 μg/mL, 100 μg/mL or 200 μg/mL, and 100 μg/mL, respectively, and stored at -20 °C as single-use aliquots. For cell treatments, cytokines were serially diluted in PBS to achieve the desired concentration, maintaining a final PBS concentration of 0.2% in cultures. Cytokines in PBS were added to bioluminescence recording media (for luminometry) or the equivalent lacking luciferin (for RT-PCR assays). For LPS-only treatments, concentrations are as indicated; for LPS/IFN-γ, 5 ng/mL of LPS and 12 ng/mL of IFN-γ were used; for IL-4 50 ng/mL was used. The RAW 264.7 recording media was prepared by dissolving powdered DMEM (Sigma-Aldrich) in Millipore-purified water (18.2 Ω resistance) to give a final concentration of 11.25 mg/mL, followed by sterile filtration using a 0.2 µm filter (Thermo Fisher Scientific), supplemented with 5% FBS, 1% HEPES (HyClone), 1% penicillin-streptomycin, 1% L-glutamine, 1% sodium pyruvate (Gibco) and 150 µg/mL D-luciferin (Pierce; not added for RT-PCR). The cytokine-containing solutions were mixed well and added to cells as designated following synchronization and removal of synchronization media. Cells were then incubated at 36.5 °C under ambient atmosphere for the duration of the respective experiment.

Conditioned media derived from EMT6 and 4T1 cells was generated as follows. Cells were plated and grown to confluence in T175 flasks using culture conditions described above. Then, growth media was removed, cells were washed with PBS, and media was replaced with conditioning media (powdered DMEM in Millipore-purified water (18.2 Ω resistance) to 13.5 g/mL, sterile filtered using a 0.2 µm filter, supplemented with 1% FBS, 1% HEPES, 1% penicillin-streptomycin, and 1% sodium pyruvate). Cells were incubated at 36.5 °C under ambient atmosphere for 72 h, after which media was removed, filtered via a 0.45 µm filter, and stored at -20 °C in single-use aliquots. Following synchronization, for experiments involving treatment with 4T1 or EMT6 conditioned media (and controls), media was removed and replaced from all samples, which received 50% RAW 264.7 bioluminescence recording media (with or without luciferin, depending on experiment), and 50% of one of the following: for non-treated (NT) control, conditioning media with 5% FBS; for FBS control, conditioning media; for 4T1 or EMT6, conditioned media from the designated breast cancer cell line. Independent of treatment, each sample had a final concentration of 1% L-glutamine and 150 µg/mL D-luciferin, which were supplemented as needed.

Conditioned media from RAW 264.7 cells was prepared similarly to above, using RAW 264.7 bioluminescence recording media (without luciferin) containing 1% FBS as conditioning media. After synchronization, for experiments involving treatment of U2OS cells with RAW 264.7 conditioned media (or controls), U2OS media was removed from all samples and replaced as follows. For NT controls, samples received 100% U2OS bioluminescence recording media (described below). Other treatments used 50% U2OS bioluminescence recording media, and 50% of one of the following: for FBS control, RAW 264.7 conditioning media; for conditioned samples, RAW 264.7 conditioned media. U2OS bioluminescence recording media was prepared by dissolving powdered DMEM (Sigma-Aldrich) in Millipore-purified water (18.2 Ωresistance) to give a final concentration of 11.25 g/mL, followed by sterile filtration using a 0.2 µm filter, supplemented with 5% FBS, 1% HEPES (HyClone), 0.25% penicillin-streptomycin, 0.35 g/L sodium bicarbonate (Fisher Scientific) and 150 µg/mL D-luciferin. Independent of treatment, each sample had a final concentration of 0.35 g/L sodium bicarbonate and 150 µg/mL D-luciferin, which were supplemented as needed.

### Bioluminescence Recording and Analysis of Time Series

Following synchronization and addition of cytokines or conditioned media, dishes were sealed with 40 mm sterile cover glass using silicon vacuum grease and subjected to a LumiCycle 32 System (Actimetrics) for monitoring at 36.5 °C for 5-7 days. Bioluminescence signals were measured every 10 min per h.

Biolumiscence recordings were pre-processed to exclude the initial 24-h transient and spikes [33]. Recordings were considered arrhythmic outliers and excluded from further analysis if their range (maximum minus minimum over time) was less than 1/3^rd^ that of the median range for that reporter and treatment. To assess the strength of rhythmicity for each rhythmic recording, a quadratic trend was removed, and then three measures were computed: the relative power of the band in the power spectral density corresponding to periods of 16h to 32h (RelPow), the rhythmicity index computed as the height of the third peak of the correlogram (RI) [34], and the maximum value of the chi square periodorgram (MaxQp) [35]. To assess circadian period and amplitude for each rhythmic recording the average of a 24-h moving window was removed, resulting in de-trended data with 12 h of data eliminated from the beginning and end of the recordings, and then the average of a 3-h moving window was used to smooth the data. The resulting time series (t=36 h to t=132 h) were fit to a damped cosine curve [33] for one estimate of the period and amplitude. Additionally, phase markers (peak, trough, mean-crossings) were identified to compute additional measures: the period as estimated by the difference in marker times from cycle to cycle, and the amplitude as measured by the difference in peak and trough heights for cycle 1 or cycle 2 [33].

## Results

### M1 and M2 polarization conditions differently alter the circadian rhythms of RAW 264.7 macrophages

To evaluate how circadian rhythms change in opposing macrophage subtypes, it is critical to generate data with sufficient resolution to facilitiate detailed analyses of oscillations. To track circadian rhythms in a time-resolved manner, with frequent sampling over the course of multiple circadian cycles, we opted to use luciferase reporters for promoter activity. Here, we stably transfected RAW 264.7 macrophage cells with a reporter for a positive (*Bmal1*) and a negative (*Per2*) component of the core circadian clock, yielding RAW 264.7-*Bmal1:luc* and RAW 264.7-*Per2:luc* cells, respectively (**Fig. S1**). LPS is an endotoxin found in outer membranes of gram negative bacteria, and results in M1 responses in RAW 264.7 macrophages [36,37]. IL-4 is secreted by T helper 2 (T_H_2) cells [16,38] and cancers [39], yielding M2 responses. To investigate circadian alterations that occur in M1 and M2 subtypes, RAW 264.7 reporter cells were polarized under the following conditions: for the M1 subtype, 5 ng/mL, 20 ng/mL, or 50 ng/mL of LPS, and a combination of 5 ng/mL of LPS and 12 ng/mL of IFN-γ (LPS/IFN-γ) were used; for the M2 subtype, 50 ng/mL of IL-4 was used [40]. To confirm cell polarization following treatments, we carried out RT-PCR to assess the presence of M1 and M2 specific markers (**Fig. S2**). As in other studies, polarization to the M1 subtype using the aforementioned conditions resulted in enhanced levels of the M1 markers *Tnf-α* (**Fig. S2A**) [36,41] and *iNos* (**Fig. S2B**) [37,41], which increased with LPS concentration and LPS/IFN-γ. Under these conditions the M2 specific marker CD206 was down-regulated. In contrast, when RAW 264.7 macrophages were polarized to the M2 subtype via IL-4, the expression of M1 markers *Tnf-α* and *iNos* decreased, as expected [40] while M2-associated CD206 increased (**Fig. S2C**).

Circadian oscillations based on *Bmal1:luc* and *Per2:luc* signals were tracked via real-time luminometry. Raw and detrended circadian data of M1 polarized cells show that average bioluminescence of *Bmal1:luc* reporters are lower (raw data shown in **Fig. 1A, S3A**; detrended data shown in **Fig. 1C, S4A**) while those of *Per2:luc* reporters vary in terms of levels (raw data shown in **Fig. 1B, S3B**; detrended data shown in **Fig. 1D, S4B**) compared to NT samples in respective RAW 264.7 reporter cell lines. In contrast, raw and detrended circadian data of M2 polarized cells show increased average bioluminescence of both *Bmal1:luc* (raw data shown in **Fig. 1A, S3A**; detrended data shown in **Fig. 1C, S4A**) and *Per2:luc* reporters (raw data shown in **Fig. 1B, S3B**; detrended data shown in **Fig. 1D, S4B**) compared to respective NT samples, in the same RAW 264.7 reporter cell lines.

**Fig. 1.**
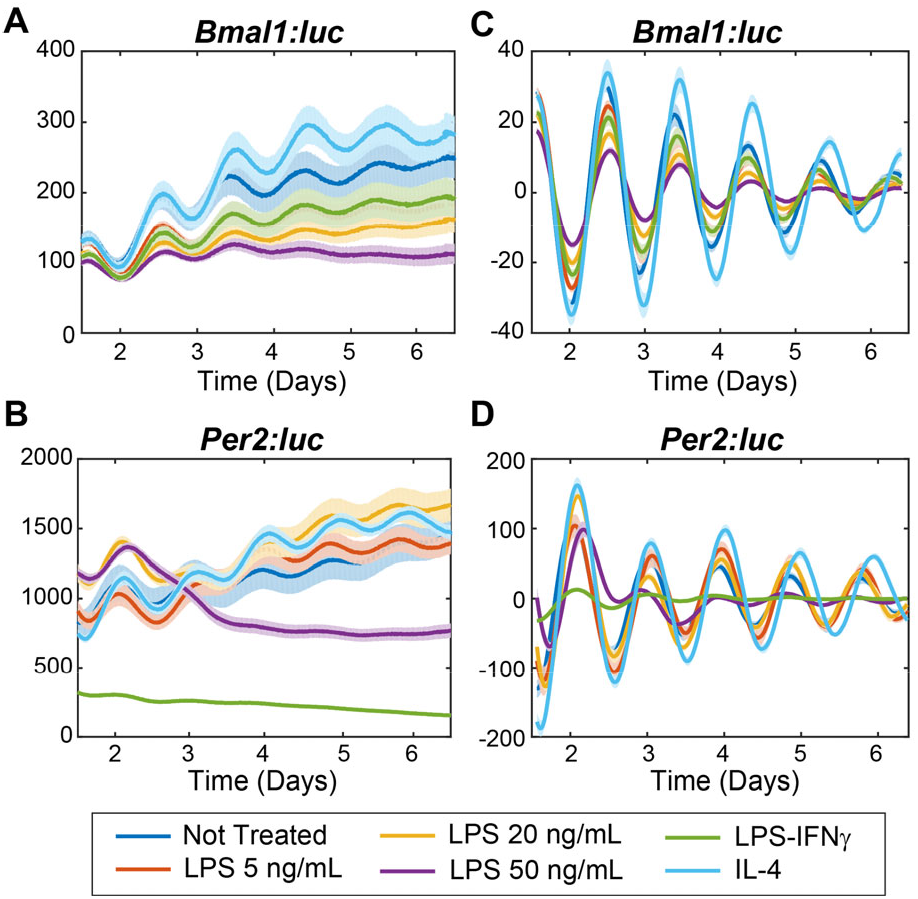
Bioluminescence rhythms of *Bmal1* and *Per2* promoter activities following cytokine treatments in RAW 264.7-*Bmal1:luc* and –*Per2:luc* cells. Shown are raw (**A**,**B**) and de-trended, smoothed (**C**,**D**) time series averaged across replicates (N=8) for each treatment. The standard error is shown with a light envelope around the mean; in some instances, this is too small to be visualized. The raw and detrended data with individual replicates can be found in **Fig. S3** and **S4**, respectively.

Circadian rhythmicity, period, and amplitude were differently altered in the M1 and M2 polarization states. The strength of rhythmicity for each time-series was assessed by three methods: the relative power in the circadian (16-32 h) band of the power spectral density (RelPow), the rhythmicity index computed as the height of the third peak of the correlogram (RI), and the maximum value of the chi-squared periodorgram (MaxQp). M1 polarization with 5 and 20 ng/mL of LPS significantly decreased *Per2:luc* rhythmicity, while 50 ng/mL of LPS reduced rhythmicity of both *Bmal1:luc* and *Per2:luc* reporters (**Fig. 2 top, S5**). The combination of LPS/IFN-γ significantly decreased rhythmicity only in the *Per2:luc* reporter (**Fig. 2B top, S5B**), compared to respective NT samples. In contrast, M2 polarization with IL-4 significantly increased the rhythmicity only in the *Per2:luc* reporter when measured using MaxQp, but not using other methods (RI, RelPow).

**Fig. 2.**
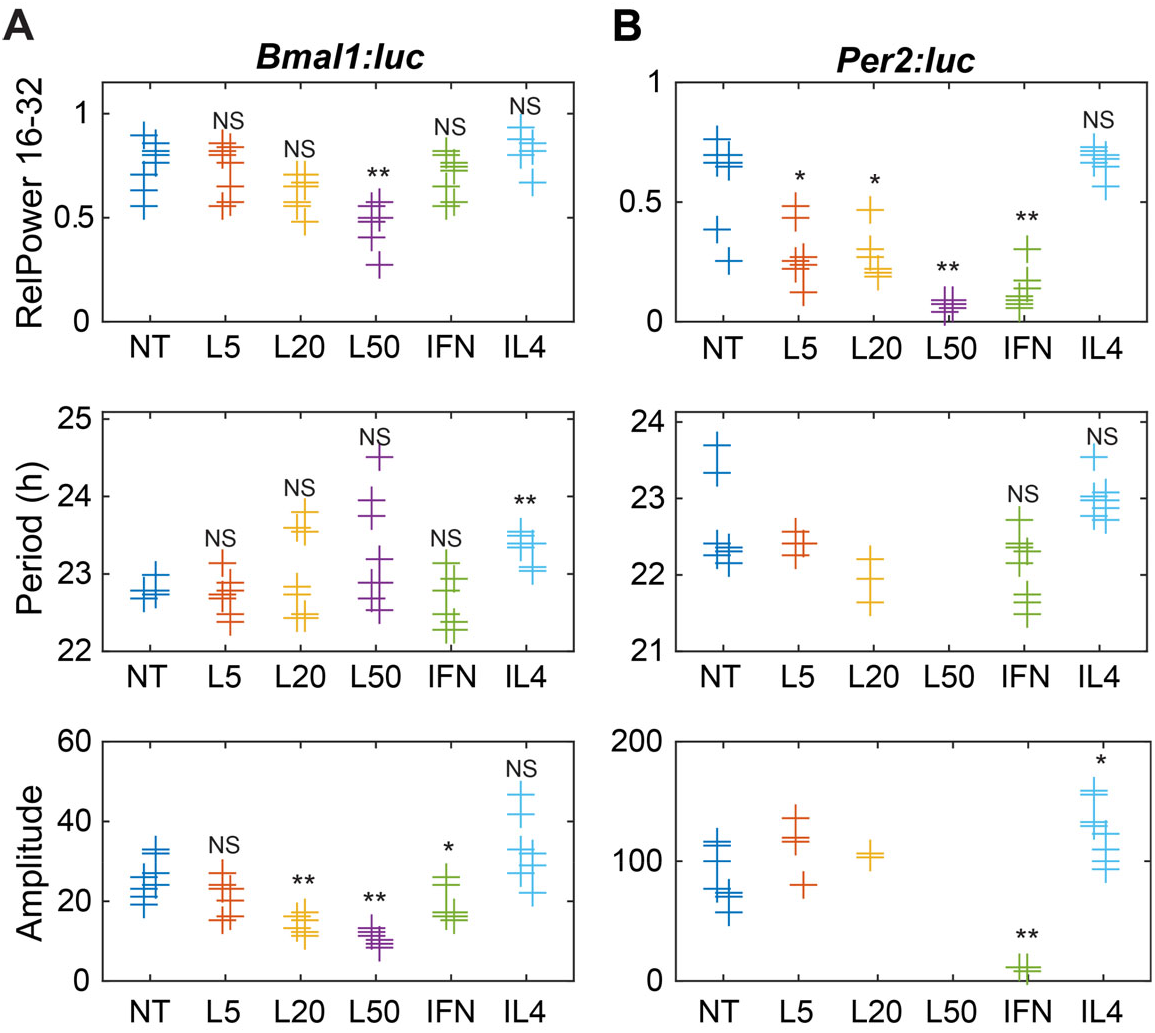
Circadian parameters for (**A**) *Bmal1* and (**B**) *Per2* oscillations following cytokine treatments. Shown are measures of strength of rhythmicity (top), period (middle), and amplitude (bottom). The measure of rhythmicity is the relative power in the 16- to 32-h band of the power spectral density. The period and amplitude are estimated by fitting each time series to a damped cosine curve. Data points are color-coded by dose; within each, data is separated by experiment (those to the left are from the first, and those to the right are from the second). The distribution of measures for each treatment is compared to that of the non-treated samples using a randomization test for difference in means (NS indicates “not significant,” ^*^ p<0.05, ^**^ p<0.01, and no indicator above a treatment indicates that there were too few data points for the test to have sufficient power). Time series that did not fit well to a damped cosine (GOF<0.9) were excluded from period and amplitude evaluations. NT = non-treated; L5 = 5 ng/mL LPS; L20 = 20 ng/mL LPS; L50 = 50 ng/mL LPS; IFN = 5 ng/mL LPS and 12 ng/mL IFN-γ.

Circadian periods were also affected by polarization. M1 polarization with LPS altered the period of RAW 264.7 cells differently depending on treatment. Decreased periods were observed following LPS treatments with 5 ng/mL (*Bmal1:luc*, mean cross up), 20 ng/mL (*Per2:luc*, trough to trough, mean CWT period), and 50 ng/mL (*Per2:luc*, mean CWT period) (**Fig. S6**). However, increased periods were observed following 50 ng/mL LPS (*Per2:luc*, mean cross up, peak to peak) (**Fig. 2, S6**). Furthermore, period values exhibited wider spreads with increasing LPS concentration in both reporters, which may be attributed to lower rhythmicity. In contrast, M2 polarization via IL-4 treatment increased periods in both reporters when measured by mean cross (up), mean cross (down), and trough to trough (**Fig. S6**), but this period enhancement was only found in the *Bmal1:luc* reporter when determined by DC Fit, peak to peak, mean CWT methods (**Fig 2A middle, S6A**). No significant period reductions were found in any of the methods used. For samples with reduced rhythmicity (time series that did not fit well to a damped cosine (GOF<0.9), no period was calculated.

Lastly, amplitudes were altered depending on polarization. M1 polarization with 20 ng/mL and 50 ng/mL of LPS, and the combination of LPS/IFN-γ significantly reduced the amplitudes of *Bmal1:luc* reporters in RAW 264.7 cells when analyzed using DC fit amplitude and peak1-trough1 (**Fig 2A bottom, S7A**), while the peak2-trough2 method showed significant period reductions for the 20 ng/mL and 50 ng/mL LPS treatments (**Fig S7A**). Amplitude reductions were also observed for LPS/IFN-γ (DC fit amplitude, peak to trough) and 50 ng/mL LPS (peak2-trough2) conditions in *Per2:luc* reporter (**Fig 2B bottom, S7B**). No significant amplitude increases were observed under any M1 polarization conditions. In contrast, amplitude enhancement was observed in IL-4 treated *Per2:luc* samples in the DC fit method (**Fig. 2B bottom, S7B**), and no significant amplitude reductions were observed under M2 polarization conditions. Where samples showed reduced rhythmicity (e.g., LPS-treated Raw 264.7-*Per2:luc*), no amplitudes were calculated.

### Exposure to breast cancer-conditioned media alters circadian rhythms of RAW 264.7 cells

Previous studies have shown that macrophages play critical roles in the dissemination of cancer cells, [42] and that cancers can polarize macrophages toward the M2 subtype [43]. These studies that imply there is a cross-talk between macrophages and the tumor microenvironment. Hence, we hypothesized that cancer cells would also affect the circadian rhythms of macrophages. To test this, first we assessed the effects of conditioned media derived from 4T1 or EMT6 mouse breast cancer cells on M1 and M2 marker levels in RAW 264.7 cells via RT-PCR. We observed that both 4T1- and EMT6-conditioned media increased the expression of the M1 markers *Tnf-α* and *iNos*, and decreased the expression of the M2 marker *CD206* (**Fig. S8**). Then, we tracked circadian rhythms of RAW 264.7 *Bmal1:luc* and *Per2:luc* reporters exposed to these conditioned media samples. Raw and detrended circadian data of both EMT6-and 4T1-conditioned media treated samples showed that average bioluminescence of *Bmal1:luc* reporters were lower (raw data shown in **Fig. 3A, S9A**; detrended data shown in **Fig. 3C, S10A**) while *Per2:luc* reporters were varying in levels (raw data shown in **Fig. 3B, S9B**; detrended data shown in **Fig. 3C, S10B**) compared to respective NT samples, in the same RAW 264.7 reporter cell lines,.

**Fig. 3.**
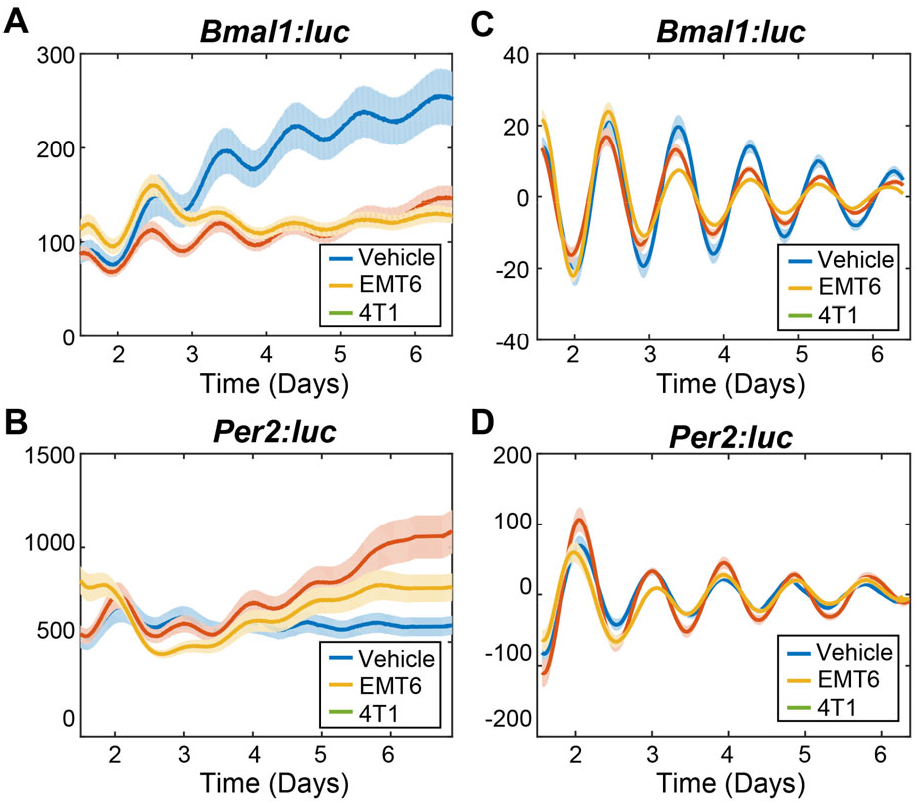
Bioluminescence rhythms of *Bmal1* and *Per2* promoter activities following macrophage exposure to cancer-conditioned media. Shown are raw (**A**,**B**) and detrended, smoothed (**C**,**D**) time series averaged across replicates (N=12) for each treatment. The standard error is shown with a light envelope around the mean; in some instances, this is too small to be visualized. The raw data and detrended data with individual replicates can be found in **Fig. S9** and **S10**, respectively.

Conditioned media derived from the highly aggressive 4T1 breast cancer cell line was found to elicit greater circadian effects than that from EMT6 cells, including with regard to reductions in circadian rhythmicity and period distributions (**Fig. 4, S11**). Rhythmicity was determined via multiple methods, all of which, except for the rhythmicity index, showed significant decreases for *Per2:luc* in 4T1-conditioned media treated cells (**Fig. 4B, S11B**). The *Bmal1:luc* reporter showed significant reduction in rhythmicity determined by the MaxQP method (**Fig. S11A**). Concurrently, treatment with EMT6 breast cancer-conditioned media resulted in no significant alterations in rhythmicity by any method used (**Fig. 4, S11**). No samples examined for any treated showed significantly increased rhythmicity. For additional circadian evaluations (e.g., period and amplitude assessments), samples that had periods outside of the 16-32 h circadian period range, or had < 0.9 damped cosine fit were considered arrhythmic and excluded.

**Fig. 4.**
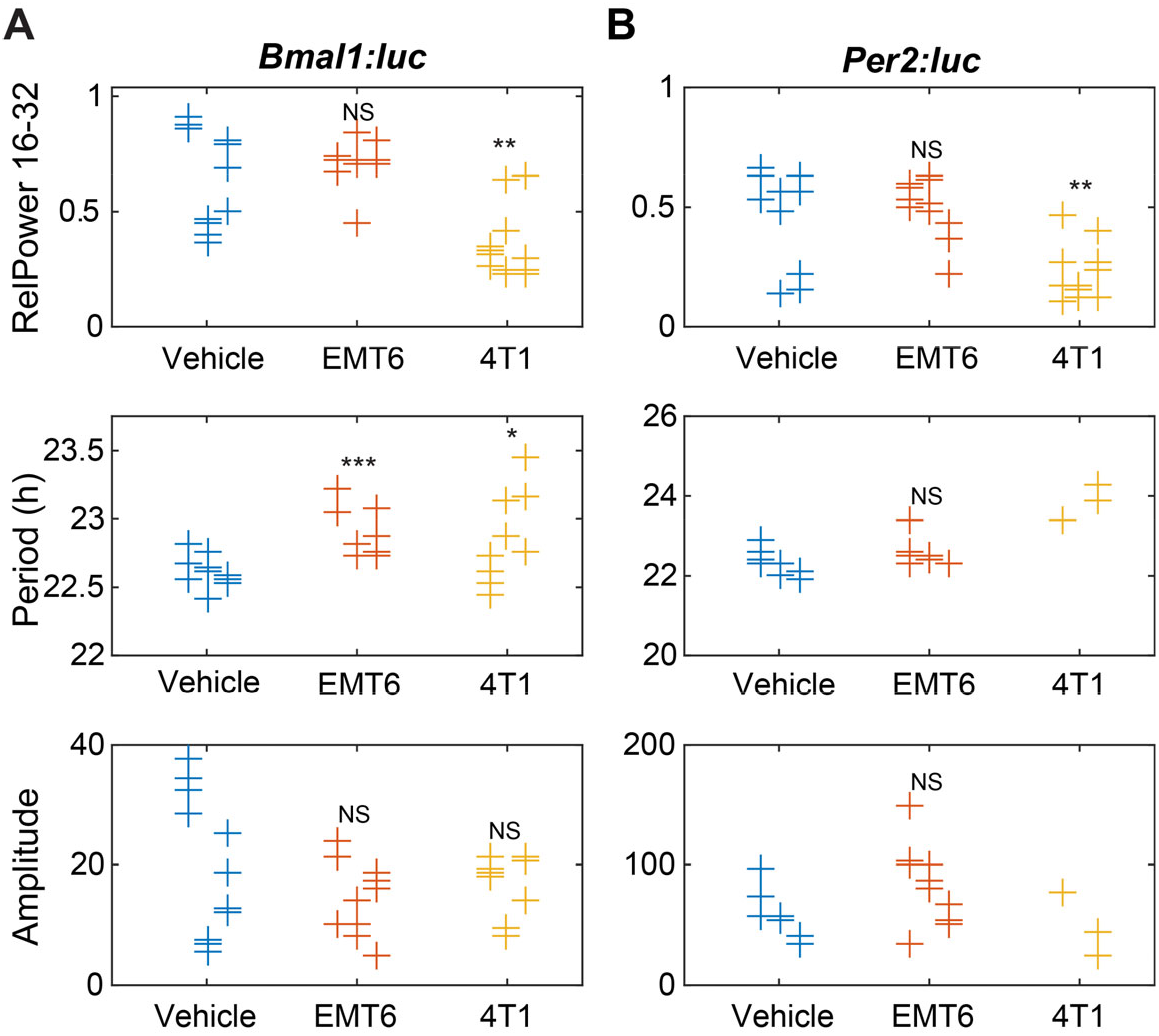
Circadian parameters for (**A**) *Bmal1* and (**B**) *Per2* oscillations following exposure to cancer cell-conditioned media. Shown are measures of strength of rhythmicity (top), period (middle), and amplitude (bottom). The measure of rhythmicity is the relative power in the 16- to 32-h band of the power spectral density. The period and amplitude are estimated by fitting each time series to a damped cosine curve. Data points are color-coded by dose; within each, data is separated by experiment (those to the left are from the first, in the middle are from the second, and to the right are from the third). The distribution of measures for each treatment is compared to that of the vehicle using a randomization test for difference in means (NS indicates “not significant”, ^*^ p<0.05, ^**^ p<0.01, and no indicator above a treatment indicates that there were too few data points for the test to have sufficient power). Time series that did not fit well to a damped cosine (GOF<0.9) were excluded from period and amplitude evaluations. (Vehicle = FBS control, EMT6 = EMT6 cell-conditioned media, 4T1 = 4T1 cell-conditioned media).

Periods were also affected by conditioned media treatments. 4T1-conditioned media treated macrophages showed significantly enhanced periods of *Bmal1:luc* and *Per2:luc. Bmal1:luc* had longer periods as determined by the mean cross up and peak to peak methods, however, it showed a period reduction using the mean cross down method (**Fig S12A**). *Per2:luc* periods were determined to increase via peak to peak, trough to trough, and mean CWT period methods; no decreases were found. EMT6-conditioned media treated samples showed significant period enhancements in the *Bmal1:luc* reporter using multiple methods (DC fit, mean cross up, peak to peak, and trough to trough; **Fig. 4A, S12A**), while the *Per2:luc* reporter showed significant period increases using peak to peak, trough to trough, and mean CWT period methods (**Fig. 4B, S12B**). No period reductions for either reporter were observed following EMT-conditioned media treatment, using any tests. It is also noteworthy that the overall distributions of period values for the 4T1-conditioned media treated samples were wider than those of controls and EMT6-conditioned media treated samples.

4T1 cell-conditioned media treated macrophage reporter cells were also affected in terms of amplitude changes, although these were not as substantial as effects on other circadian characteristics. *Bmal1:luc* oscillations showed significantly reduced amplitude using the peak2-trough2 method (**Fig. S13A**); however, the other approaches used did not result in statistically significant changes. No amplitude evaluations showed effects on the *Per2:luc* reporter, or either *Bmal1:luc* or *Per2:luc* signals for cells treated with EMT6-conditioned media. (**Fig. 4B, S13**).

### RAW 264.7-conditioned media affects the circadian rhythms of U2OS cells

Macrophages have shown different responses and interactions with cancers. In some cases, macrophages facilitate the metastasis and invasion of breast cancer cells [44,45] while in others, they reduce cell proliferation [46,47]. Here, we investigated whether and how conditioned media derived from naïve macrophages affects osteosarcoma cells, which have diminished growth following macrophage exposure, and their circadian outputs. We used the human osteosarcoma cell line U2OS, which is a well-established model for circadian studies [48–50] to evaluate the effects of macrophage-conditioned media on circadian oscillations of cancer cells. While the cell lines are derived from different species, interactions between the two have been previously confirmed [51]. We used U2OS cells, stably transfected with *Bmal1:luc* and *Per2:luc* luciferase reporters [31], and treated them with conditioned media harvested from naïve RAW 264.7 cells. The circadian oscillations of U2OS-*Bmal1:luc* and – *Per2:luc* were subsequently tracked via luminometer for 5-7 days.

Raw and detrended circadian data of RAW 264.7-conditioned media treated U2OS cells showed that the average bioluminescence levels of *Bmal1:luc* reporters were lower (raw data shown in **Fig. 5A, S14A**; detrended data shown in **Fig. 5C, S15A**) while *Per2:luc* reporter bioluminescence levels were higher (raw data shown in **Fig. 5B, S14B**; detrended data shown in **Fig. 5C, S15B**), compared to respective NT samples, in the same U2OS reporter cell line.

**Fig. 5.**
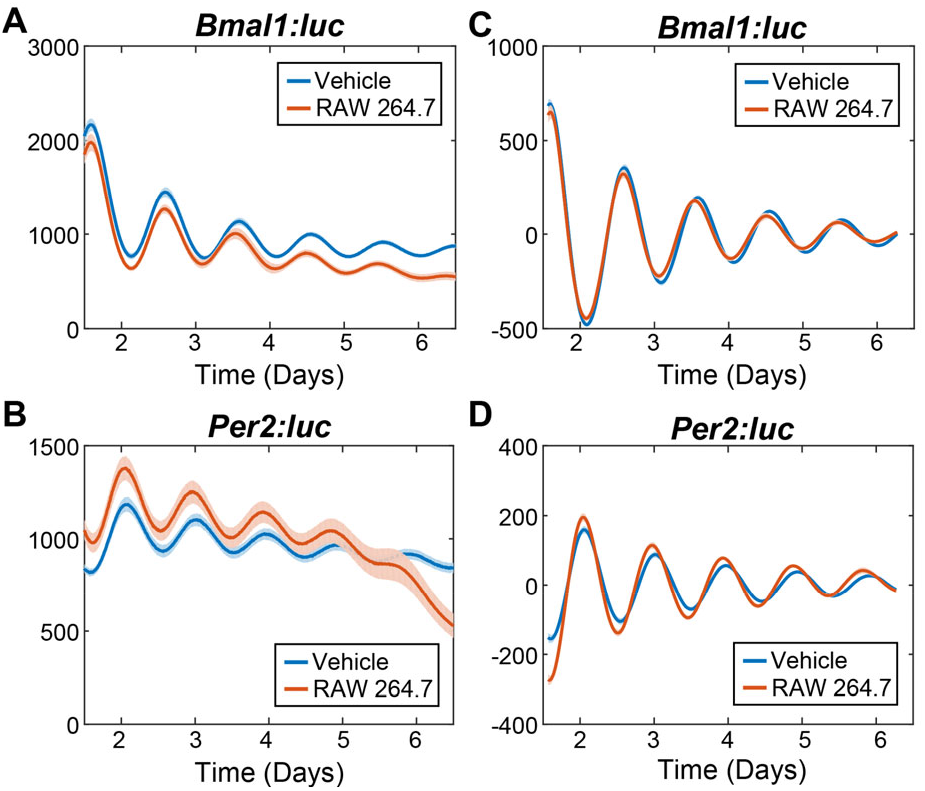
Bioluminescence rhythms of *Bmal1* and *Per2* promoter activities in U2OS cells following exposure to conditioned media from macrophages. Shown are raw (**A**,**B**) and de-trended, smoothed (**C**,**D**) time series averaged across replicates (N=8) for each treatment. The standard error is shown with a light envelope around the mean; in some instances, this is too small to be visualized. The raw and detrended data with individual replicates can be found in **Fig. S14** and **S15**, respectively.

The *Per2:luc* reporter showed significantly decreased rhythmicity in the RelPow 16-32 h and rhythmicity index methods while *Bmal1:luc* reporter did not show any significant rhythmicity alterations (**Fig 6, S16**). No samples showed increased rhythmicity. Period was also altered due to the conditioned media treatment. *Per2:luc* reporter showed significantly decreased period in the DC fit, mean cross up, peak to peak methods while *Bmal1:luc* reporter did not show any significant period alterations (**Fig 6, S17**). No samples showed increased period. Overall, the conditioned media treated samples showed higher distribution of period values compared to NT samples. Furthermore, *Per2:luc* reporter showed significantly increased amplitude in the DC fit mean damped amplitude, and peak2-trough2 methods while *Bmal1:luc* reporter did not show any significant amplitude alterations (**Fig 6, S18**). No samples showed decreased amplitude. In follow-up experiments, we also evaluated U2OS cell treatments with conditioned media from M1- and M2-polarized macrophages, however the results obtained did not differ from those presented here and thus are not shown.

**Fig. 6.**
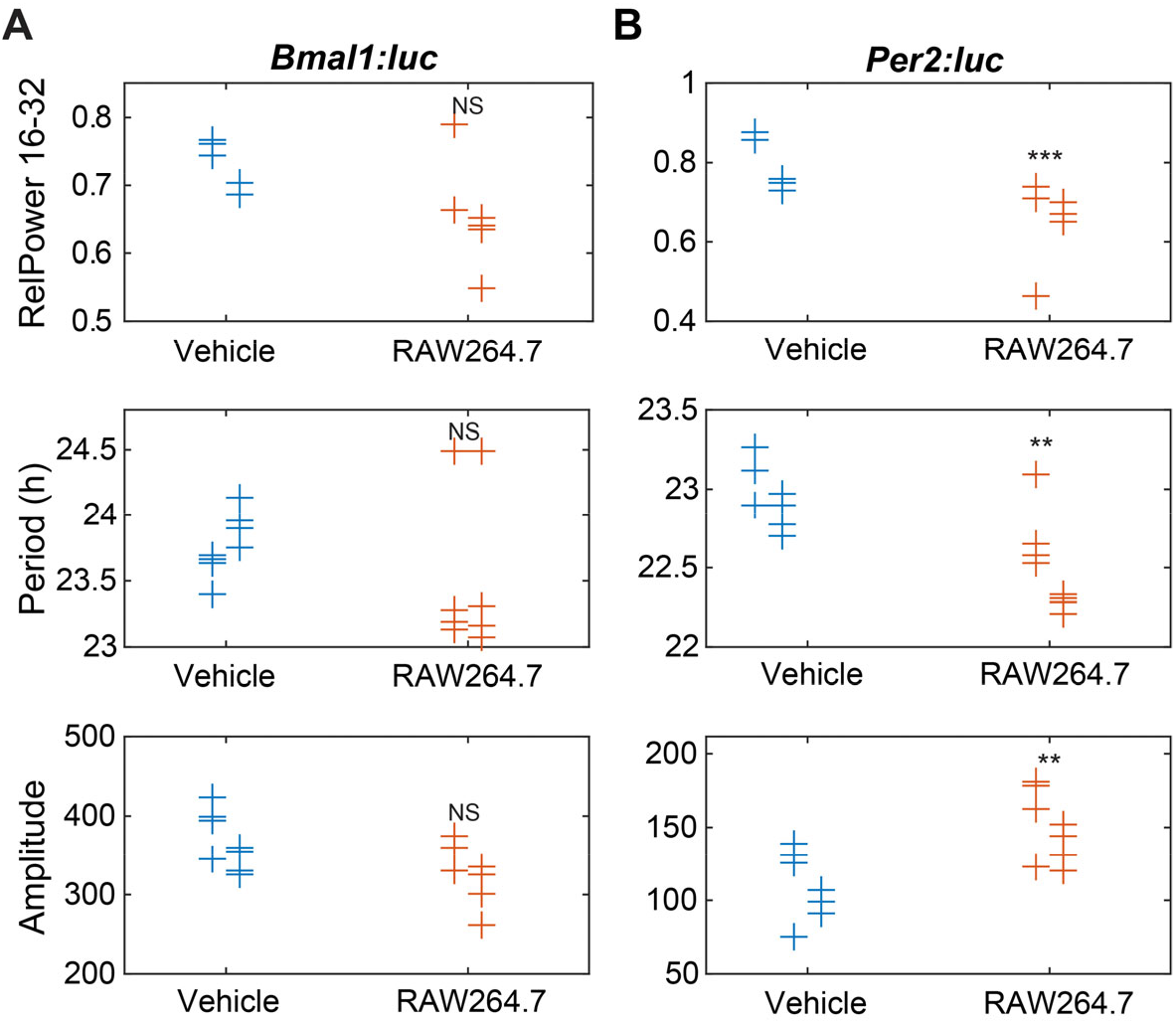
Circadian parameters of (**A**) *Bmal1* and (**B**) *Per2* in U2OS cells following exposure to macrophage conditioned media. Shown are measures of strength of rhythmicity (top), period (middle), and amplitude (bottom). The measure of rhythmicity is the relative power in the 16- to 32-h band of the power spectral density. The period and amplitude are estimated by fitting each time series to a damped cosine curve. Data points are color-coded by dose; within each, data is separated by experiment (those to the left are from the first, in the middle are from the second, and to the right are from the third). The distribution of measures for each treatment is compared to that of the vehicle using a randomization test for difference in means (NS indicates “not significant”, ^***^ p<0.001, ^**^ p<0.01, and no indicator above a treatment indicates that there were too few data points for the test to have sufficient power). Time series that did not fit well to a damped cosine (GOF<0.9) were excluded from period and amplitude evaluations. (Vehicle = FBS control, RAW264.7 = RAW 264.7 cell-conditioned media).

## Discussion

Macrophages have diverse functions that range from pro-inflammatory (e.g., generation of reactive oxygen and nitrogen species) to immune-suppressing (e.g., tissue remodeling and angiogenesis) activities. To produce the appropriate responses to physiological or environmental challenges, macrophages are plastic -- they can be polarized to broadly M1 (pro-inflammatory) or M2 (anti-inflammatory) subtypes [15,16]. Previous studies have shown that 8-15% of the macrophage transcriptome, including genes involved in pathogen recognition and responses, has circadian oscillations [17,19]. Multiple functions and characteristics of macrophages have also been shown to occur in circadian manner, such as recruitment to infected tissue [8], generation of chemokines and cytokines, and phagocytosis [20,21]. Based on the distinct, opposing functions of the two general macrophage subtypes, and macrophages’ inherent associations with circadian rhythms, we hypothesized that the oscillations of circadian genes are differently altered depending on stimuli.

Here, we studied the relationship between macrophage polarization and core clock oscillations by tracking the signals of *Bmal1:luc* and *Per2:luc* reporters in RAW 264.7 cells exposed to various stimuli, using real-time luminometry. We performed two series of experiments in this regard: first, we used cytokines to polarize cells to standard immune-stimulating (M1) and –suppressing (M2) states; then, we exposed macrophages to conditioned media from two different murine breast cancer cell types, 4T1 (more aggressive) and EMT6 (less aggressive). We found that opposing polarizations of macrophages differentially altered their circadian rhythms, and that the circadian alterations observed following conditioned media treatments are similar to those of M1 polarized macrophages.

It has been previously shown that mRNA/protein levels of core circadian clock genes in macrophages are altered under pro-inflammatory conditions [25,27]. LPS treatment significantly reduced *Bmal1* mRNA levels in peritoneal macrophages [25], mouse BMDM and peritoneal macrophages, and human macrophages and peripheral blood mononuclear cells (PBMC) [27]. In our own study, we similarly saw that LPS-treated *Bmal1*:*luc* reporters have lower bioluminescence levels compared to NT samples in both raw (**Fig. 1A, S3A**) as well as detrended (**Fig. 1B, S4A**) data, suggesting that M1 polarization with LPS weakens *Bmal1* circadian rhythms. It has also been shown that treatment with a high concentration of LPS significantly increased *Per2* mRNA levels in mouse BMDMs [27]. However, a luminometry study carried out with *mPer2:luc* mouse-derived peritoneal macrophages showed that 5 ng/mL LPS reduced, but 20 ng/mL and 100 ng/mL LPS increased non-detrended signal intensities for a portion of the experiment (24-54 hours) compared to control [25], while another study showed that 50 ng/mL LPS reduced non-detrended rhythms in mouse BMDMs over four days [26]. Likewise, our data showed (over 24-72 hours) that 5 ng/mL reduced raw bioluminescence rhythms of *Per2:luc* in RAW 264.7 cells, but (over 24 hours to 1 week) 20 ng/mL and (over 24-72 hours) 50ng/mL LPS resulted in increases (**Fig. 1B**, and **S3B**). However, after ∼3 days, we saw increased raw bioluminescence levels for 5 ng/mL LPS and reduced bioluminescence for 50 ng/mL LPS-treated samples compared to NT.

For LPS/IFN-γ-treated samples, reduced bioluminescence was observed, compared to NT, for the entire time-series (**Fig. 1B**, and **S3B**), as seen previously with *mPer2:luc* BMDM [26]. Our data showed that circadian amplitudes were reduced when cells were exposed to LPS or LPS/ IFN-γ (**Fig. 2**, and **S7**), which were also similar to previous studies showing that LPS or LPS/IFN-γ treatments can reduce the amplitudes of peritoneal macrophages [25] and BMDMs [26] in *mPer2:luc* mice. While LPS or LPS/ IFN-γ treatments showed no significant period effects in *mPer2:luc* BMDM [26], and in our study (**Fig. 2** and **S6**), LPS treatment has been shown to induce a subtle lengthening of period in *mPer2:luc* peritoneal macrophages [25]. We also observed that M1 polarization with 5 and 20 ng/mL of LPS, and combination of LPS/IFN-γ significantly decreased *Per2:luc* rhythmicity, while 50 ng/mL of LPS reduced rhythmicity of both *Bmal1:luc* and *Per2:luc* reporters (**Fig. 2, S5**). However, other studies have not assessed cellular rhythmicity.

Weakened circadian amplitudes observed in the M1 subtype may be caused by the generation of reactive oxygen species (ROS) in these macrophages [25]. M1 polarization increases expression of ROS producing genes (e.g., *Tnfα* and *iNos*). While we did not find any changes to viability, high concentrations of ROS have been shown to kill M1 macrophages [24,54], and less viable cells could have damped/lower amplitude rhythms. Under M2 polarization (described further below), such ROS generating genes are down-regulated (**Fig. S2**), and enhanced amplitudes were observed (**Fig. 1, 2**). However, another likely reason for decreased *Bmal1* oscillations is the production of proinflammatory miRNA-155 in response to LPS treatment. Curtis et al. showed that miR-155 levels are inversely correlated with those of *Bmal1*, which possesses two miR-155 binding sites in its promoter, and is unaffected following LPS treatment in the presence of a miR-155 antagomir [27].

While the circadian effects of M2 polarized macrophages have not been as thoroughly studied, M2 polarization/IL-4 (20 ng/mL) treatment has been shown to decrease mRNA and protein levels of REV-ERBα (a *Bmal1* transcriptional repressor) in differentiated THP-1 macrophages [29]. Such reduction of REV-ERBα should increase levels of *Bmal1*; we also found increased levels of raw bioluminescence for the *Bmal1:luc* reporter (**Fig. 2a**) upon M2 polarization in our study. The increased Bmal1 should also drive *Per2* expression, which we also observed via amplitude enhancement in *Per2:luc* (**Fig. 2b, S7**), and was found by Chen et al. [26]. However, while that study did not find any effects on period, we observed increased periods in both reporters following M2 polarization (**Fig. 2, S6**). Furthermore, we also found that M2 polarization significantly increased rhythmicity of the *Per2:luc* reporter (**Fig. 2, S5**).

There is significant evidence to indicate cross-talk between macrophages and tumor microenvironments, but the contributions of macrophages also depends on cancer type [42,43,55,56]. Similarly, macrophage responses to conditioned media can vary. For example, conditioned media derived from high tumor grade MDA-MB-231 cells (human breast cancer) has shown to upregulate both M1 and M2 markers of human CD14^+^ macrophages [43], while murine BMDMs exposed to 4T1 breast cancer conditioned media, and splenocytes cocultured with 4T1 cells expressed higher levels of M1 marker genes including *Tnf-α* [55,56]. However, the effects of tumor microenvironments on macrophage circadian rhythms are unknown. To evaluate the interactions between the two, we treated RAW 264.7-*Bmal1:luc* and -*Per2:luc* macrophages with EMT6 (less aggressive) and 4T1 (highly aggressive) murine mammary carcinoma [57] derived-conditioned media and analyzed their circadian effects. Our data showed that the more aggressive 4T1 breast cancer-conditioned media reduced macrophage rhythmicity akin to M1 polarization conditions, which was not surprising given the similar marker profile as determined by RT-PCR (**Fig. S8**). On the other hand, the less aggressive EMT6-derived conditioned media did not affect rhythmicity, but instead yielded significantly altered periods. Altogether our data suggests that mouse breast cancer conditioned media polarizes macrophages M1 subtype as a defense mechanism to protect the host against the cancer.

As mentioned above, macrophages can result in positive or negative effects following interaction with cancers. Previous studies show that macrophages promote metastasis and invasion of breast cancer cells [44,45], while others show that macrophages inhibit osteosarcoma cell growth via inhibition of cell proliferation and increased phagocytosis [46,47,51]. Considering the previously investigated effects of macrophage-conditioned media on cancer cells, and the interactions between macrophages and osteosarcomas, we wished to evaluate whether the former might also influence circadian rhythms of the latter. We treated human osteosarcoma reporter cell lines, U2OS-*Bmal1:luc* and -*Per2:luc* with conditioned media harvested from naïve RAW 264.7 cells. We observed reduction in rhythmicity and amplitude enhancement of the U2OS oscillations. As circadian rhythms are found to be increasingly disrupted with disease severity and oncogenic characteristics [32] and even subtle renormalization of disrupted rhythms can reduce oncogenic features [48], our data highlights another connection between circadian oscillations and cancer, which should be studied further.

Taken together, we show that the circadian rhythms of macrophages are influenced by their polarization states. While M1 polarization was associated with significant loss of rhythmicity, M2 polarization resulted in amplitude and period enhancements. We also found that macrophage circadian effects translate to cultures with cancer cell-conditioned media, where M1-like marker characteristics were accompanied by M1-like circadian oscilations. Finally, we show that conditioned media of macrophages enhances the circadian rhythms of another cancer cell type (U2OS), which macrophages act against in oncogenic environments.

This raises a hypothesis that macrophages can alter the circadian rhythms of tumors and cancer cell types to affect their oncogenic features. In the future, this relationship should be examined with other cancer models, including to determine whether macrophages negatively affect the circadian oscillations of cancer cells/types with which they interact to facilitate disease. In terms of macrophages themselves, it is of interest to study whether the lack of rhythmicity in inflammatory macrophages is due to loss of synchrony by performing single-cell experiments, and to assess whether this circadian divergence confers benefits in terms of immune response and host defense.

## Acknowledgements

J. A. M.-R. was supported by a fellowship from the Chemistry-Biology Interface Training Program (National Research Service Award (T32 GM008515)) from the National Institutes of Health. H.-H. L. was supported by a University of Massachusetts Amherst Chemistry-Biology Interface (CBI) training fellowship. We would like to thank Tanya Leise (Mathematics & Statistics, Amherst College) for helpful discussions regarding analyses.

