## Supporting Information for "Macrophage Circadian Rhythms are Differentially Affected Based on Stimuli"

Luciferase Assay

Cells were seeded in 24-well plates with 500 µL cells at a density of 2 x $10^5$ cells/mL. Once the cells reached confluence (approximately 48 h), they were lysed and assessed using the Luciferase Assay System (Promega) according to the manufacturer's instructions. Luminescence from the RAW 264.7-*Bmal1:luc* and *-Per2:luc* cells was measured via a SpectraMax M5 multi-mode microplate reader.

RT-PCR Experiments

Each condition used for RT-PCR experiments was performed in biological triplicates. Following treatment of cells with cytokines or conditioned media and incubation for 48 h, RNA was harvested as described previously.[1] Briefly, RNA was harvested from cells using a PureLink RNA Mini Kit (Ambion) according to the manufacturer's instructions. Each RNA sample was reverse transcribed to cDNA using 50 µM random hexamers (Applied Biosystems), 10 mM dNTPs (Thermo Scientific), 40 U/µL RNaseOut (Invitrogen), 200 U/µL SuperScript IV Reverse Transcriptase (Invitrogen), 100 mM DTT (Invitrogen), and 5x Super Script IV buffer (Invitrogen). RT-PCR was performed in 96-well plates using a CFX Connect Real-Time System (Bio-Rad). Each reaction consisted of 100 ng cDNA, 10 µL iTaq universal SYBR Green Supermix (Bio-Rad), 4 µM each forward and reverse primer (Integrated DNA Technologies), and RNAse-free water (Fisher) to 20 µL. The samples were centrifuged and processed using an initial denaturation at 95 °C for 3 min, followed by 40 cycles of 95 °C denaturation for 10 s, and 58 °C annealing/extension for 30 s. The primers used were *β-actin* forward (5'- GAT CAG CAA GCA GGA GTA CGA -3'), reverse (5'- AAA ACG CAG CGC AGT AAC AGT -3'); *iNos* forward (5'- GTT CTC AGC CCA ACA ATA CAA GA -3'), reverse (5'- GTG GAC GGG TCG ATG TCA C -3'); *Tnf-α* forward (5'- CCT GTA GCC CAC GTC GTA G -3'), reverse (5'- GGG AGT AGA CAA GGT ACA ACC C -3'); *CD206* forward (5'- GGA TGT TGA TGG CTA CTG GA -3'), reverse (5'- AGT AGC AGG GAT TTC GTC TG -3'). Relative *Tnf-α, iNos, and CD206* expression levels were determined by comparing $C_t$ values for these genes to *β-actin* (control) via $2^{\Delta\Delta C_t}$ method.[2] Three biological replicates with three technical replicates for each were analyzed for each condition.

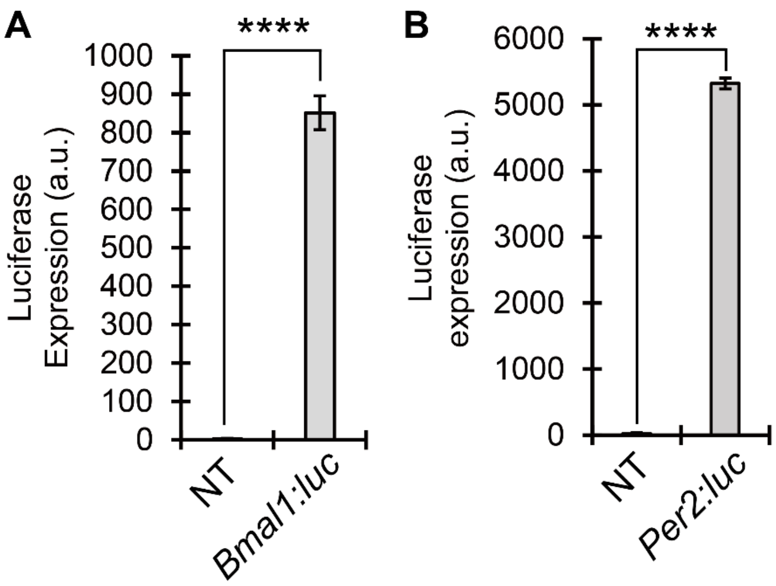


**Fig. S1** Luciferase assay data obtained for the stable transfections of (**A**) *Bmal1:luc* and (**B**) *Per2:luc* into RAW 264.7 cells. Each condition has three technical replicates (n=3). Paired student T-tests were used to calculate significance between conditions (***p<0.0001). The error bars represent standard deviation of the mean. NT = non-transfected, parental RAW 264.7 cells.

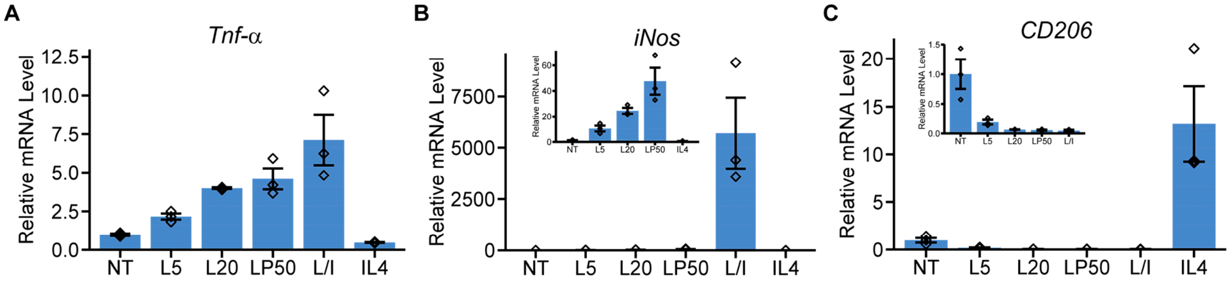


**Fig. S2** Relative mRNA levels of *Tnf-α*, *iNos*, and *CD206* following polarization via cytokine treatments in RAW 264.7 cells. mRNA levels were quantified using RT-qPCR. As expected, LPS and LPS/IFN-γ treatments upregulated M1 markers (**A**) *Tnf-α* and (**B**) *iNos*, and downregulated the M2 marker (**C**) *CD206*. IL-4 treatment downregulated (**A**) *Tnf-α* and (**B**) *iNos*, and upregulated (**C**) *CD206*. Insets within **B** and **C** show data without L/I and IL4, respectively. Each treatment contained three biological replicates, with three technical replicates each. Each mean is shown as a diamond; error bars represent standard error of the mean (SEM). NT = non-treated; L5 = 5 ng/mL LPS; L50 = 50 ng/mL LPS; L/I= 5 ng/mL LPS and 12 ng/mL IFN-γ; IL4 = 50 ng/mL IL-4.

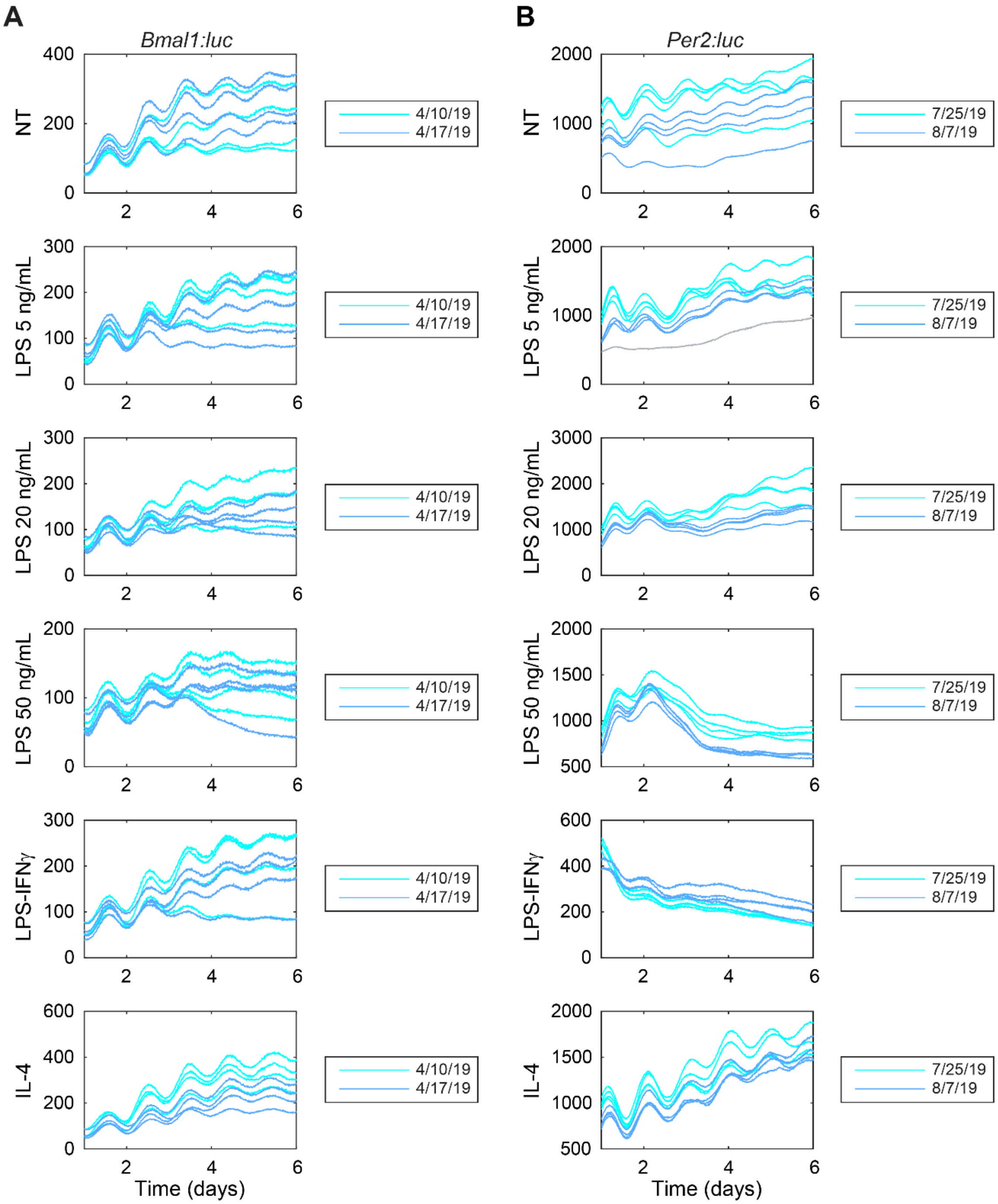


**Fig. S3** Raw bioluminescence following cytokine treatments. Shown are time series for (**A**) *Bmal1* and (**B**) *Per2* promoter activities. Each time series is color-coded by experiment date (N=4 each). There is one outlier (gray, a *Per2:luc* recording from LPS 5 ng/mL treatment), that when de-trended has a range smaller than one-third of the median range for the given reporter and treatment.

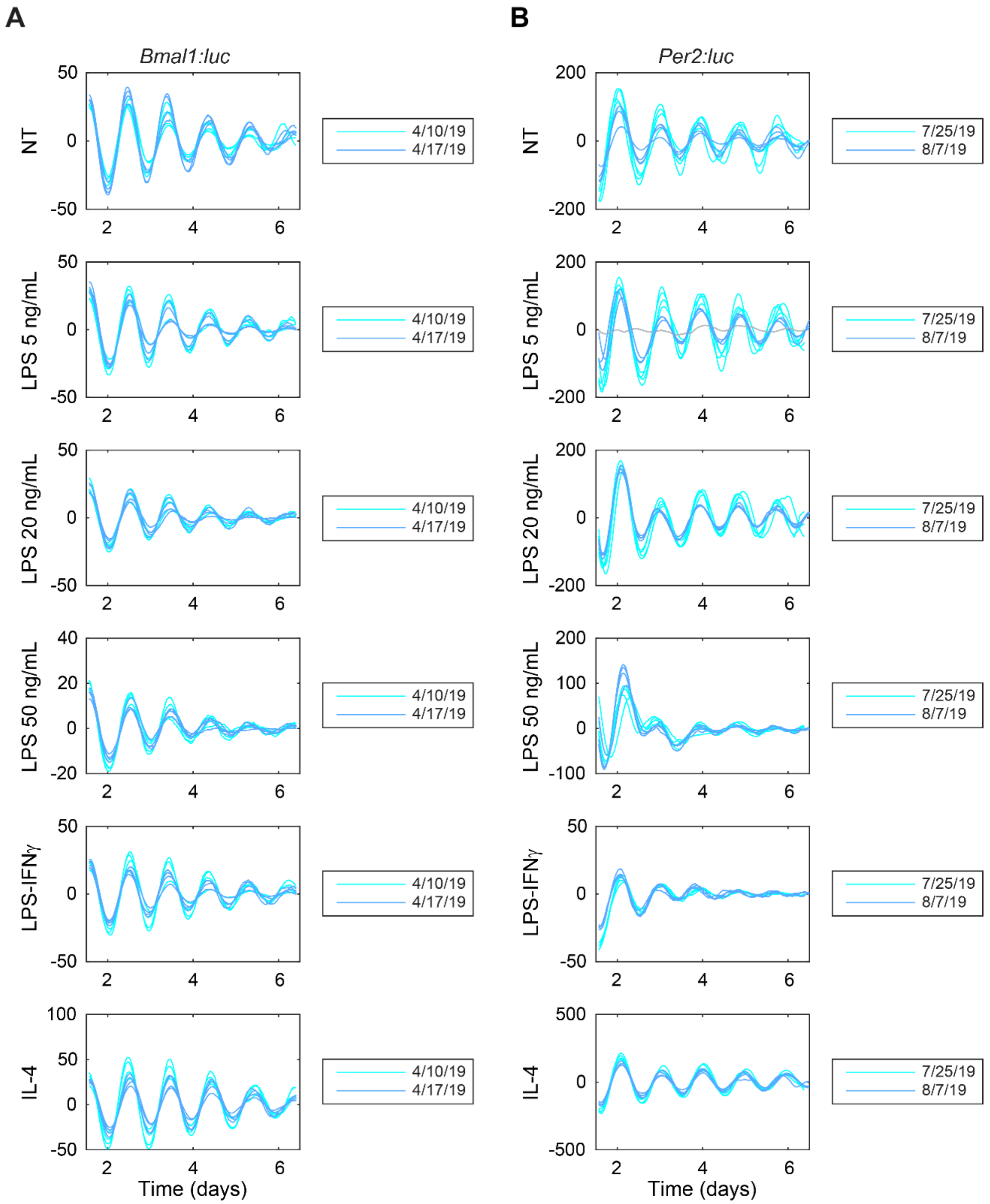


**Fig. S4** Detrended bioluminescence following cytokine treatments. Shown are time series that have been de-trended by subtracting the mean of a 24-h sliding window and smoothed with the mean a 3-h sliding window for (**A**) *Bmal1* and (**B**) *Per2* promoter activities. Each time series is color-coded by experiment date (N=4 each). There is one outlier (gray, a *Per2:luc* recording from 5 ng/mL LPS treatment), that when detrended has a range smaller than one-third of the median range for the given reporter and treatment.

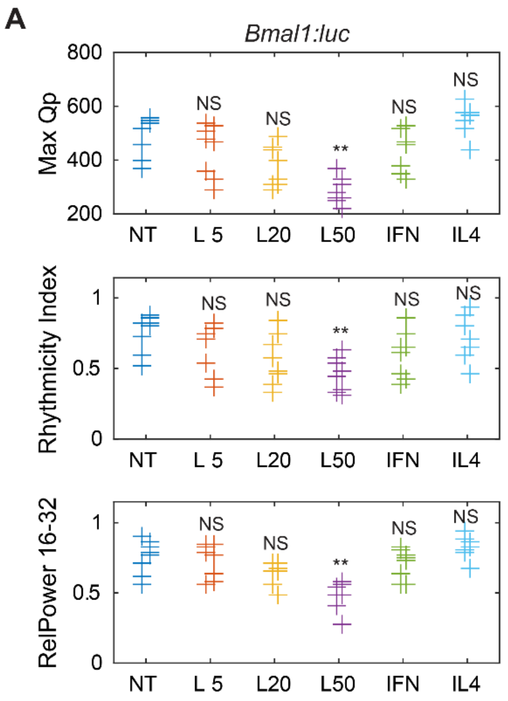


**Fig. S5** Comparison of results from application of multiple measures of rhythmicity for cytokine-treated cells. Shown are data from three analyses: Maximum Qp value from Chi-Square Periodogram (top), Rhythmicity Index from a correlogram (middle), and the relative power in the 16- to 32-h band of the power spectral density (bottom; also shown in **Fig. 2** and duplicated here for comparison) for (**A**) *Bmal1* and (**B**) *Per2* promoter activities. Data points are color-coded by dose; within each, data is separated by experiment (those to the left are from the first, and those to the right are from the second). The distribution of measures for each treatment is compared to that of the non-treated samples using a randomization test for difference in means (NS indicates "not significant", * $p<0.05$, ** $p<0.01$, and no indicator above a treatment indicates that there were too few data points for the test to have sufficient power). NT=non-treated; L5 = 5 ng/mL LPS; L20 = 20 ng/mL LPS; L50 = 50 ng/mL LPS; IFN = 5 ng/mL LPS and 12 ng/mL IFN-γ.

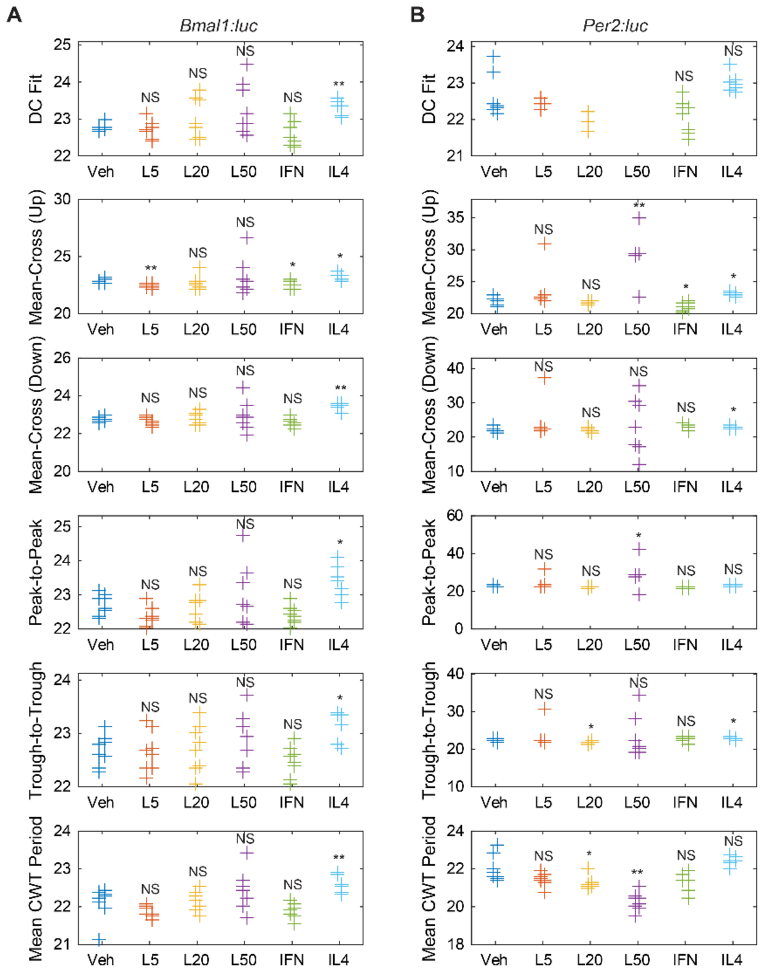


**Fig. S6** Comparison of results from application of multiple types of measures for period determination for cytokine-treated cells. Shown are results from six methods for estimating period: damped cosine fit (DC Fit, top; also shown in **Fig. 2** and reproduced here for comparison), average time between mean-crossings as bioluminescence rises (Mean-Cross (Up), second row), average time between mean-crossings as bioluminescence falls (Mean-Cross (Down), third row), average time between peaks (Peak-to-Peak, fourth row), average time between troughs (Trough-to-Trough, fifth row), and the continuous wavelet-estimate averaged across time (Mean CWT Period, bottom) for (**A**) *Bmal1* and (**B**) *Per2* promoter activities. Data points are color-coded by dose; within each, data is separated by experiment (those to the left are from the first, and those to the right are from the second). The distribution of measures for each treatment is compared to that of the vehicle (non-treated samples) using a randomization test for difference in means (NS indicates "not significant", * $p<0.05$, ** $p<0.01$, and no indicator above a treatment indicates that there were too few data points for the test to have sufficient power). Veh=Vehicle; L5 = 5 ng/mL LPS; L20 = 20 ng/mL LPS; L50 = 50 ng/mL LPS; IFN = 5 ng/mL LPS and 12 ng/mL IFN-γ.

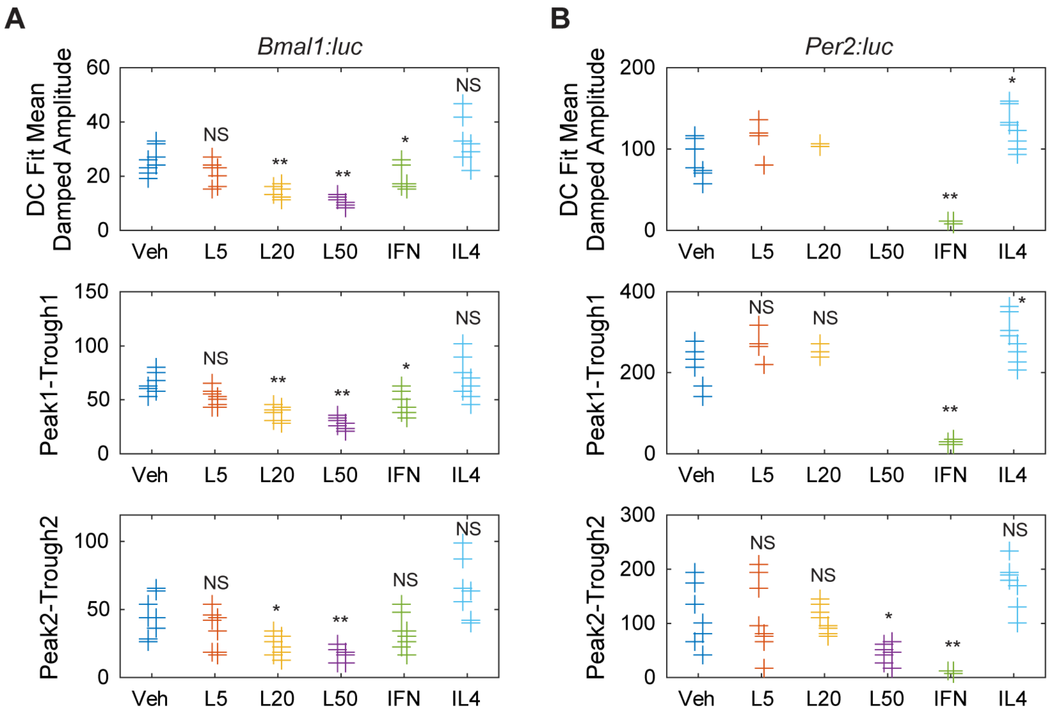


**Fig. S7** Comparison of results from application of multiple types of measures for amplitude determination for cytokine-treated cells. Shown are results from three methods for estimating period: damped cosine fit (top; also shown in **Fig. 2**, and reproduced here for comparison), peak-to-trough amplitude of first cycle (middle), and peak-to-trough amplitude of second cycle (bottom) for (**A**) *Bmal1* and (**B**) *Per2* promoter activities. Data points are color-coded by dose; within each, data is separated by experiment (those to the left are from the first, and those to the right are from the second). The distribution of measures for each treatment is compared to that of the vehicle (non-treated samples) using a randomization test for difference in means (NS indicates "not significant", * $p<0.05$, ** $p<0.01$, and no indicator above a treatment indicates that there were too few data points for the test to have enough power). Veh = Vehicle, L5 = 5 ng/mL LPS; L20 = 20 ng/mL LPS; L50 = 50 ng/mL LPS; IFN = 5 ng/mL LPS and 12 ng/mL IFN-γ.

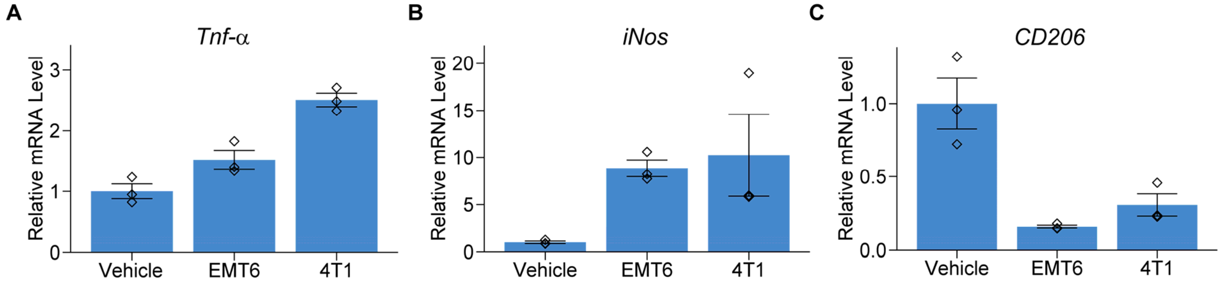


**Fig. S8** Relative mRNA levels of *Tnf-α*, *iNos*, and *CD206* in RAW 264.7 cells following exposure to cancer cell-conditioned media. mRNA levels were quantified using RT-qPCR. EMT6 and 4T1 conditioned media treatments resulted in increased levels of (**A**) TNF-α and (**B**) iNOS, and decreased (**C**) CD206. Each treatment contained three biological replicates, with three technical replicates each (whose means are shown as diamonds). Error bars represent standard error of the mean (SEM). Vehicle = FBS control, EMT6 = EMT6 cell-conditioned media, 4T1 = 4T1 cell-conditioned media.

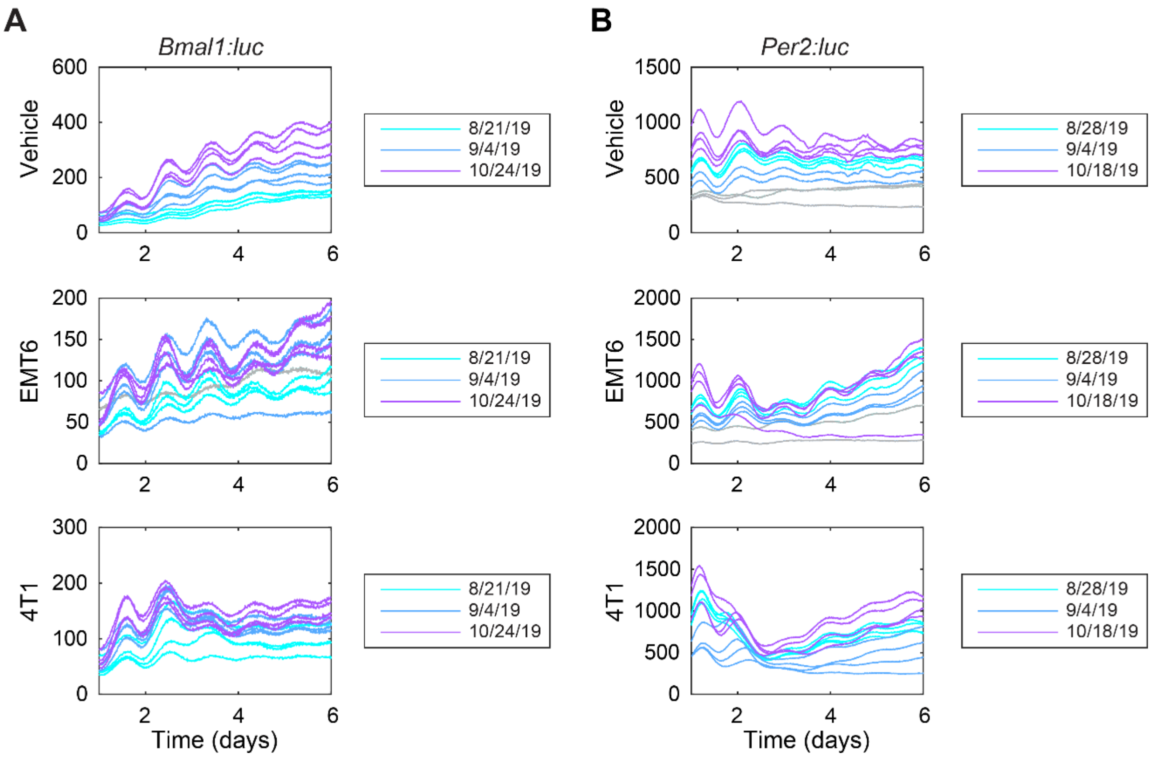


**Fig. S9** Raw bioluminescence following conditioned media treatments. Shown are time series for (**A**) *Bmal1* and (**B**) *Per2* promoter activities. Each time series is color-coded by experiment date (N=4 per experiment). Outliers (gray) are time series that have a range smaller than 1/3rd of the median range for the given reporter and treatment (for *Bmal1:luc* recordings, N=1 from EMT6 cell-conditioned media; for *Per2:luc* recordings, N=3 from FBS control and N=2 from EMT6 cell-conditioned media).

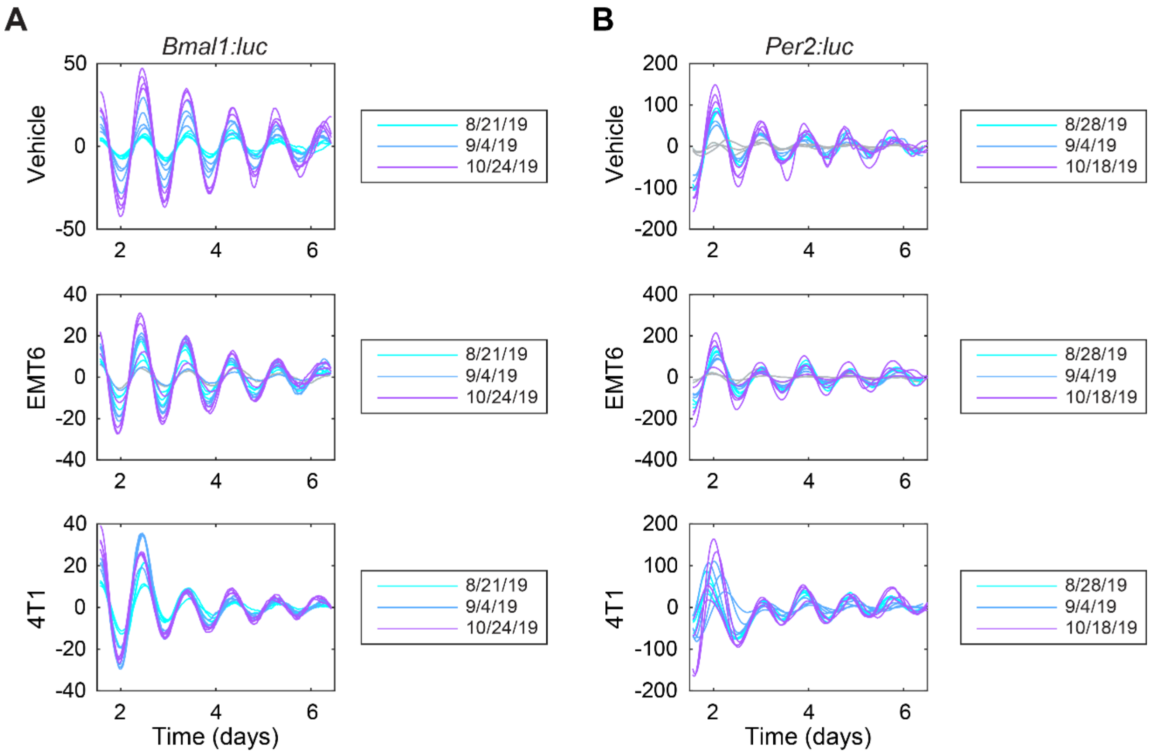


**Fig. S10** Detrended bioluminescence following conditioned media treatments. Shown are time series that have been de-trended by subtracting the mean of a 24-h sliding window and smoothed with the mean a 3-h sliding window for (**A**) *Bmal1* and (**B**) *Per2* promoter activities. Each time series is color-coded by experiment date (N=4 each). Outliers (gray) are time series that have a range smaller than 1/3rd of the median range for the given reporter and treatment (for *Bmal1:luc* recordings, N=1 from EMT6 cell-conditioned media; for *Per2:luc* recordings, N=3 from FBS control and N=2 from EMT6 cell-conditioned media).

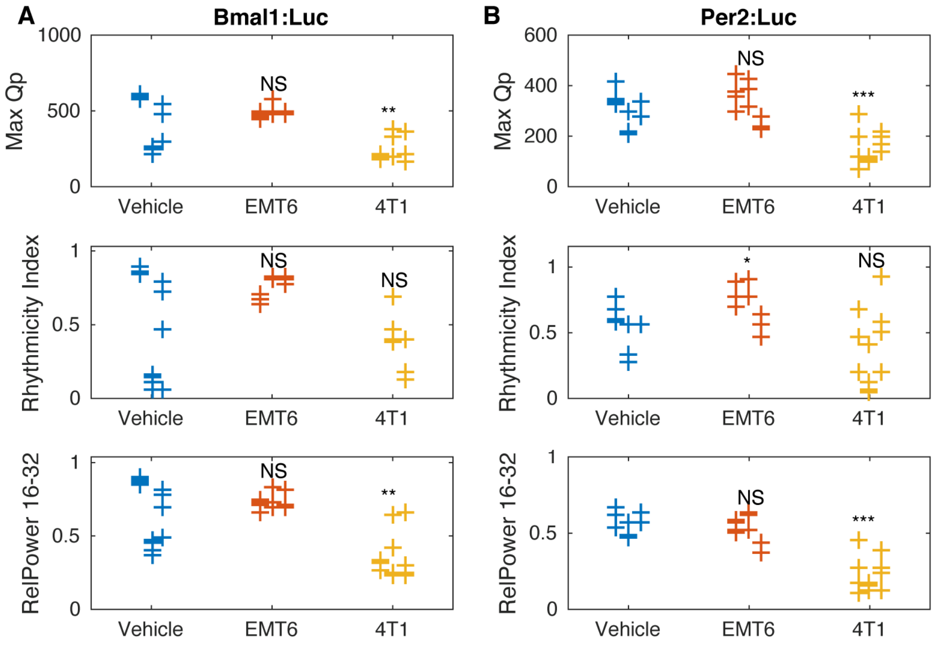


**Fig. S11** Comparison of results from application of multiple measures of rhythmicity following exposure to conditioned media. Shown are data from three analyses: Maximum Qp value from Chi-Square Periodogram (top), Rhythmicity Index from a correlogram (middle), and the relative power in the 16- to 32-h band of the power spectral density (bottom; also shown in **Fig. 4** and duplicated here for comparison) for (**A**) *Bmal1* and (**B**) *Per2* promoter activities. Data points are color-coded by dose; within each, data is separated by experiment (those to the left are from the first, in the middle are from the second, and to the right are from the third). The distribution of measures for each treatment is compared to that of the vehicle using a randomization test for difference in means (NS indicates "not significant", * $p<0.05$, ** $p<0.01$, and no indicator above a treatment indicates too few data points for the test to have enough power). (Vehicle = FBS control, EMT6 = EMT6 cell-conditioned media, 4T1 = 4T1 cell-conditioned media).

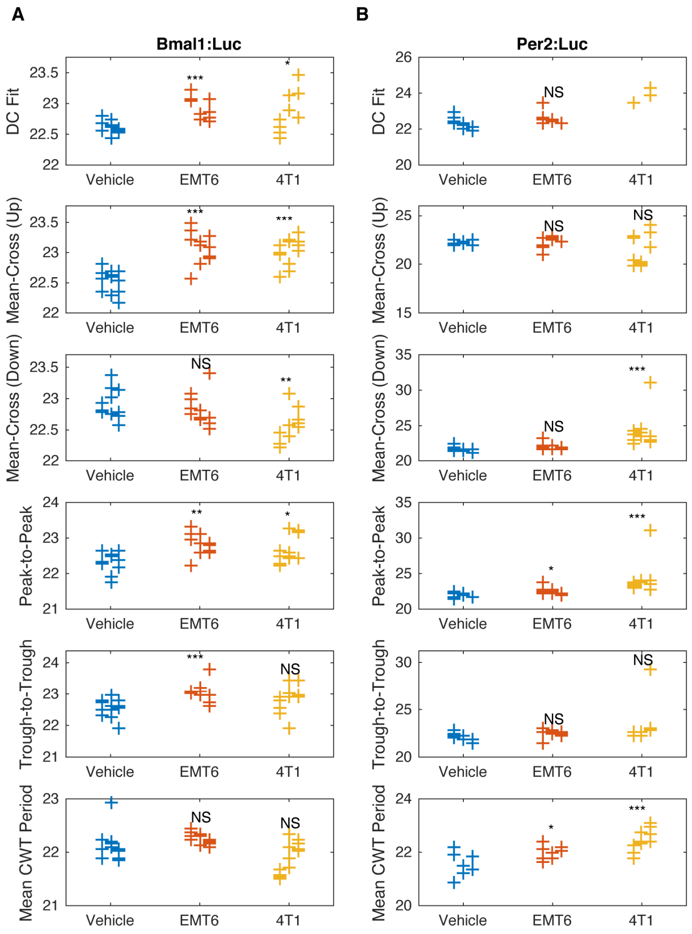


**Fig. S12** Comparison of results from application of multiple types of measures for period determination following conditioned media treatments. Shown are results from six methods for estimating period: damped cosine fit (DC Fit Mean, top; also shown in **Fig. 4**), average time between mean-crossings as bioluminescence rises (Mean-Cross (Up), second row), average time between mean-crossings as bioluminescence falls (Mean-Cross (Down), third row), average time between peaks (Peak-to-Peak, fourth row), average time between troughs (Trough-to-Trough, fifth row), and the continuous wavelet-estimate averaged across time (Mean CWT Period, bottom) for (**A**) *Bmal1* and (**B**) *Per2* promoter activities. Data points are color-coded by dose; within each, data is separated by experiment (those to the left are from the first, in the middle are from the second, and and to the right are from the third). The distribution of measures for each treatment is compared to that of the vehicle using a randomization test for difference in means (NS indicates "not significant", * p<0.05, ** p<0.01, and no indicator above a treatment indicates that there were too few data points for the test to have sufficient power). (Vehicle = FBS control, EMT6 = EMT6 cell-conditioned media, 4T1 = 4T1 cell-conditioned media).

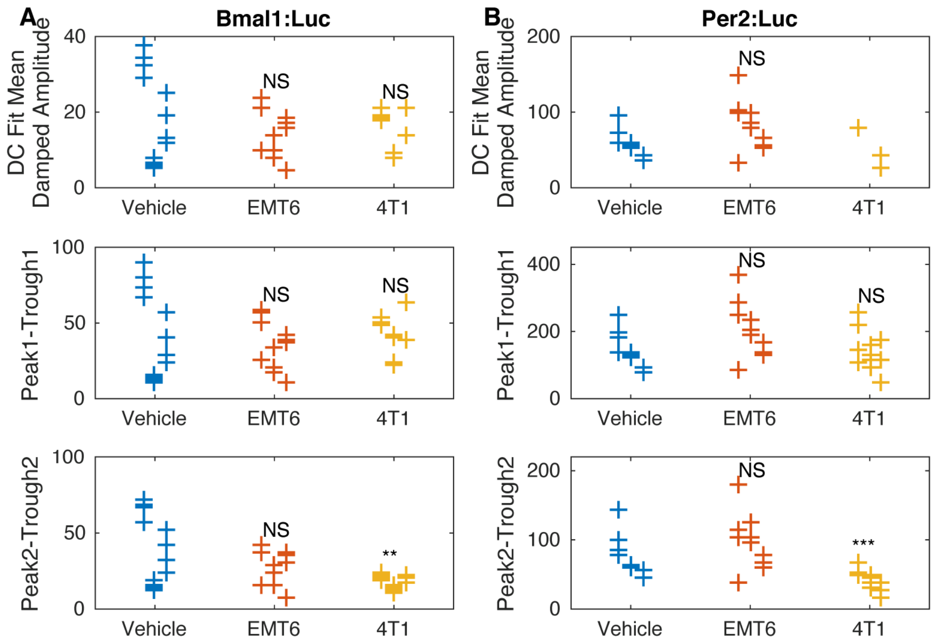


**Fig. S13** Comparison of results from application of multiple types of measures for amplitude determination following exposure to cancer cell-conditioned media. Shown are results from three methods for estimating amplitude: damped cosine fit (top; also shown in **Fig. 4**, and reproduced here for comparison), peak-to-trough amplitude of first cycle (middle), and peak-to-trough amplitude of second cycle (bottom) for (**A**) *Bmal1* and (**B**) *Per2* promoter activities. Data points are color-coded by dose; within each, data is separated by experiment (those to the left are from the first, in the middle are from the second, and to the right are from the third). The distribution of measures for each treatment is compared to that of the vehicle using a randomization test for difference in means (NS indicates "not significant", * p<0.05, ** p<0.01, and no indicator above a treatment indicates that there were too few data points for the test to have sufficient power). (Vehicle = FBS control , EMT6 = EMT6 cell-conditioned media, 4T1 = 4T1 cell-conditioned media).

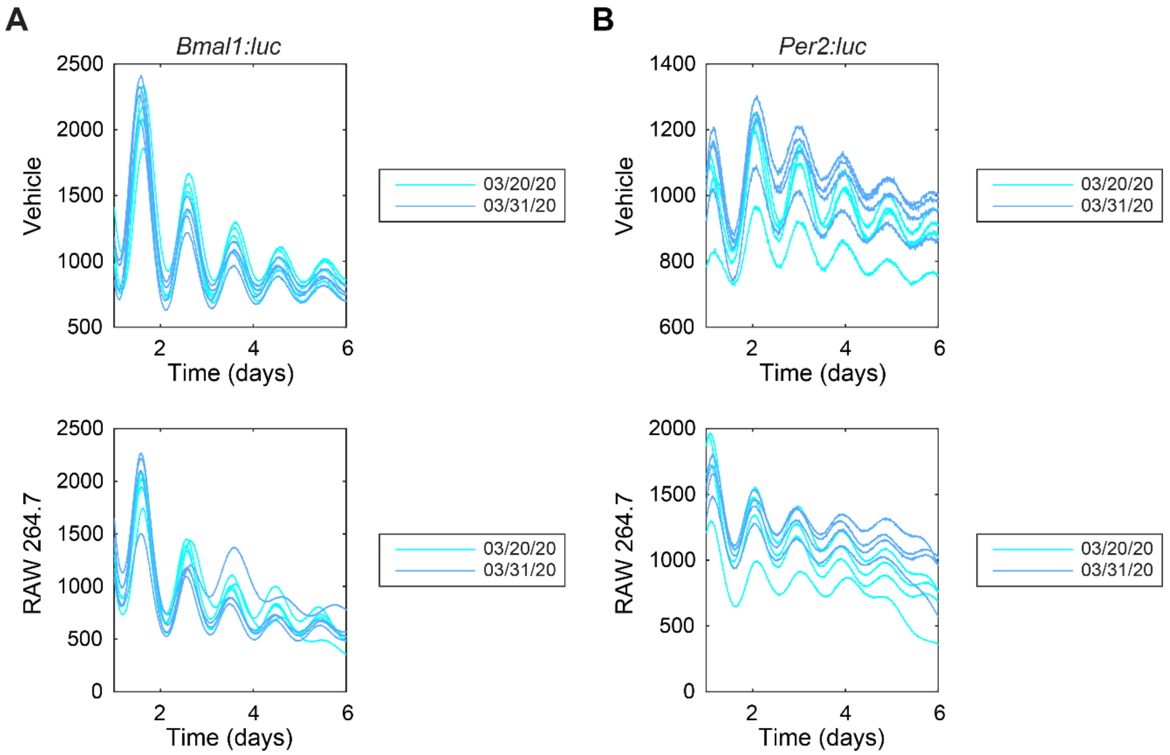


**Fig. S14** Raw bioluminescence following conditioned media treatments in U2OS cells. Shown are time series for (**A**) *Bmal1* and (**B**) *Per2* promoter activities. Each time series is color-coded by experiment date (N=4 each).

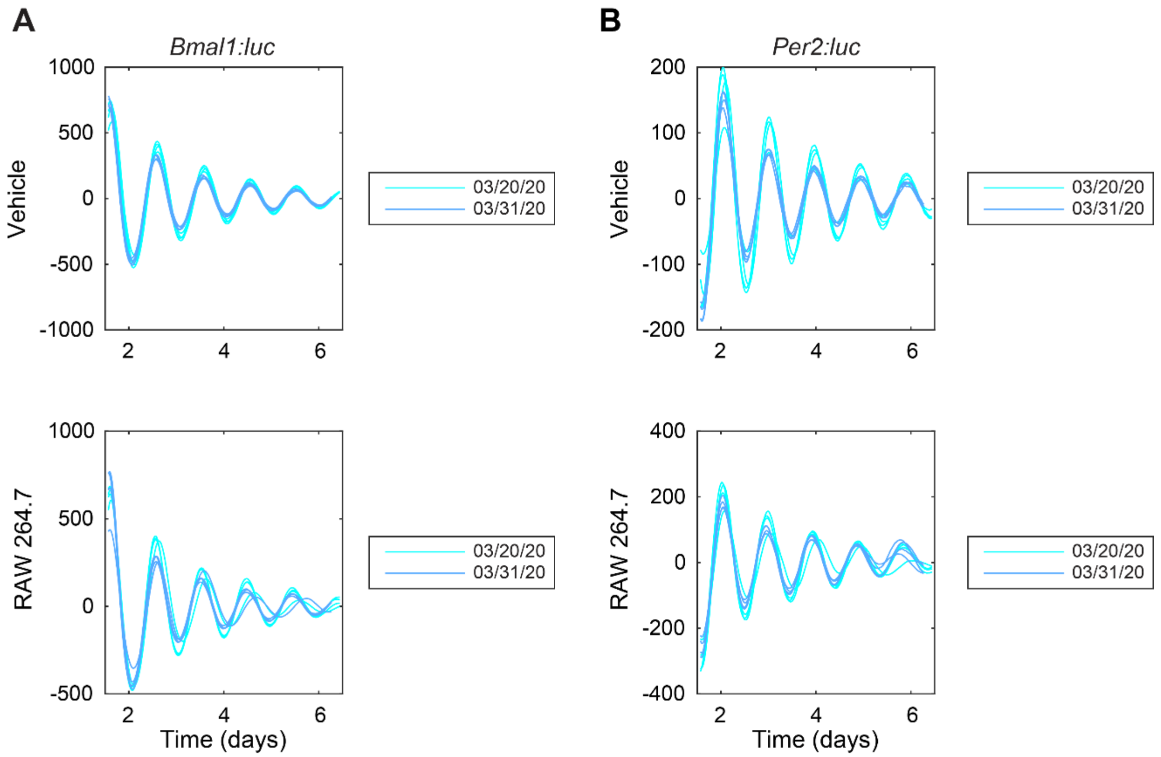


**Fig. S15** De-trended bioluminescence following conditioned media treatments in U2OS cells. Shown are time series that have been de-trended by subtracting the mean of a 24-h sliding window and smoothed with the mean a 3-h sliding window for (**A**) *Bmal1* and (**B**) *Per2* promoter activities. Each time series is color-coded by experiment date (N=4 each).

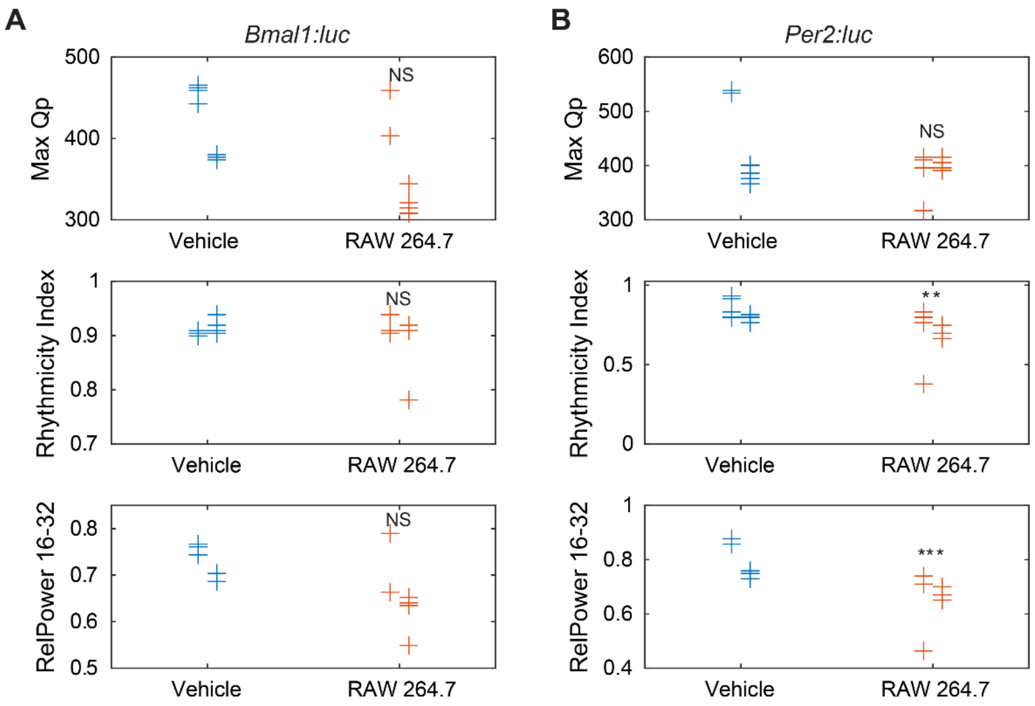


**Fig. S16** Comparison of results from application of multiple measures of rhythmicity following U2OS cell exposure to macrophage-conditioned media.. Shown are data from three analyses: Maximum Qp value from Chi-Square Periodogram (top), Rhythmicity Index from a correlogram (middle), and the relative power in the 16- to 32-h band of the power spectral density (bottom; also shown in **Fig. 6** and duplicated here for comparison) for (**A**) *Bmal1* and (**B**) *Per2* promoter activities. Data points are color-coded by dose; within each, data is separated by experiment (those to the left are from the first, and to the right are from the second. The distribution of measures for each treatment is compared to that of the vehicle using a randomization test for difference in means (NS indicates "not significant", *** p<0.001, **p<0.01, and no indicator above a treatment indicates too few data points for the test to have enough power). (Vehicle = FBS control, RAW 264.7 = RAW 264.7 cell-conditioned media).

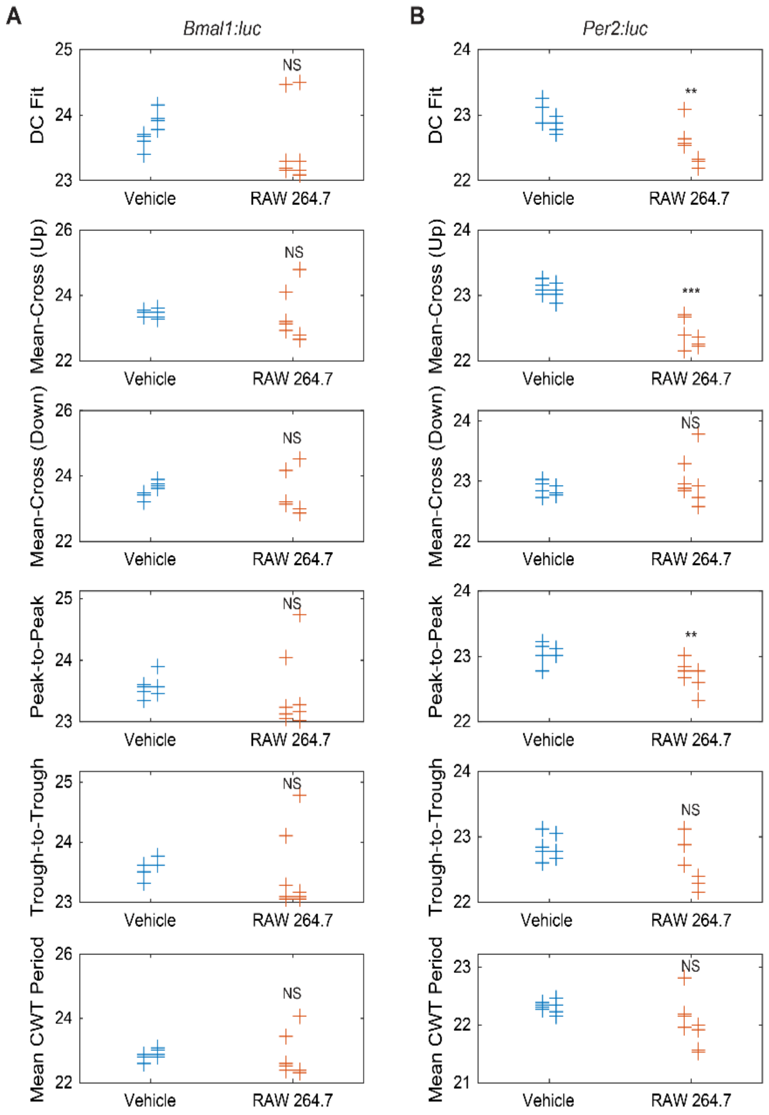


**Fig. S17** Comparison of results from application of multiple types of measures for period determination following U2OS cell exposure to macrophage-conditioned media. Shown are results from six methods for estimating period: damped cosine fit (DC Fit Mean, top; also shown in **Fig. 6** and duplicated here for comparison), average time between mean-crossings as bioluminescence rises (Mean-Cross (Up), second row), average time between mean-crossings as bioluminescence falls (Mean-Cross (Down), third row), average time between peaks (Peak-to-Peak, fourth row), average time between troughs (Trough-to-Trough, fifth row), and the continuous wavelet-estimate averaged across time (Mean CWT Period, bottom) for (**A**) *Bmal1* and (**B**) *Per2* promoter activities. Data points are color-coded by dose; within each, data is separated by experiment (those to the left are from the first and to the right are from the second). The distribution of measures for each treatment is compared to that of the vehicle using a randomization test for difference in means (NS indicates “not significant”, *** $p<0.001$, ** $p<0.01$, and no indicator above a treatment indicates that there were too few data points for the test to have sufficient power). (Vehicle = FBS control, RAW 264.7 = RAW 264.7 cell-conditioned media).

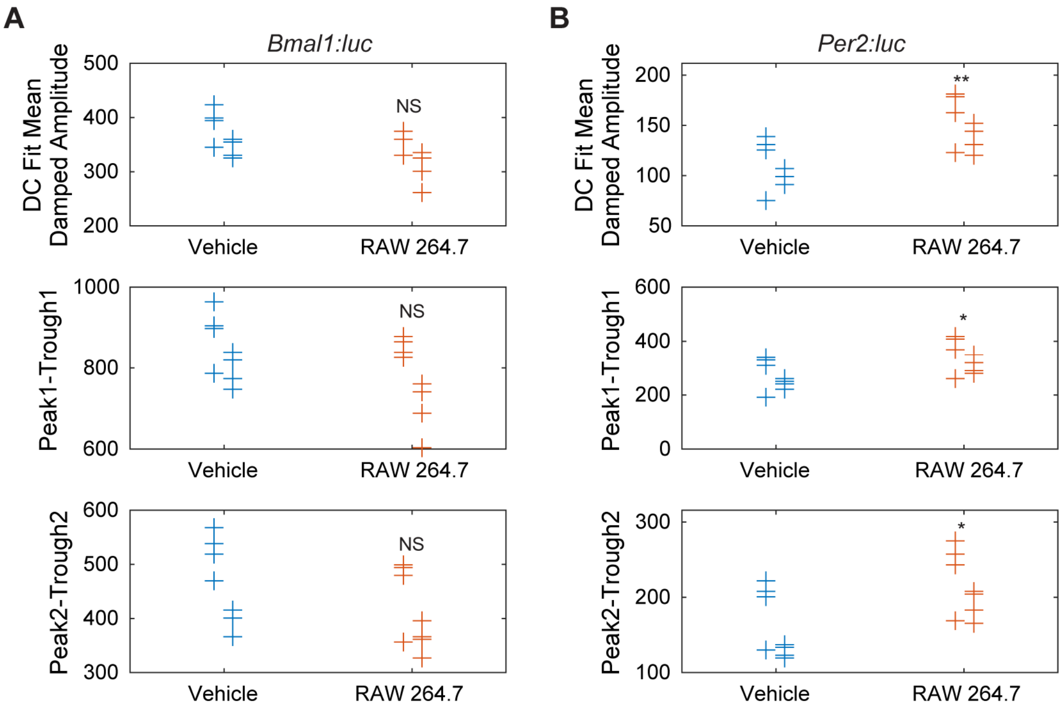


**Fig. S18** Comparison of results from application of multiple types of measures for amplitude determination following U2OS cell exposure to macrophage-conditioned media. Shown are results from three methods for estimating amplitude: damped cosine fit (top; also shown in **Fig. 6**, and reproduced here for comparison), peak-to-trough amplitude of first cycle (middle), and peak-to-trough amplitude of second cycle (bottom) for (**A**) *Bmal1* and (**B**) *Per2* promoter activities. Data points are color-coded by dose; within each, data is separated by experiment (those to the left are from the first, and to the right are from the third. The distribution of measures for each treatment is compared to that of the vehicle using a randomization test for difference in means (NS indicates "not significant", * $p<0.05$, ** $p<0.01$, and no indicator above a treatment indicates that there were too few data points for the test to have sufficient power). (Vehicle = FBS control, RAW 264.7 = RAW 264.7 cell-conditioned media).
